# Genomic Determinants of Phage Activity Against *Pseudomonas aeruginosa*: Roles of Receptors, Defence Systems, and Anti-Defences

**DOI:** 10.64898/2026.02.22.706922

**Authors:** Andrew Vaitekenas, Chris J. Malajczuk, Liza Mantjani, Phoebe G. Carr, Joshua J. Iszatt, Renee N. Ng, Samuel T. Montgomery, Yuliya Karpievitch, Stephen M. Stick, Anthony Kicic, PhageWA

**Affiliations:** Wal-Yan Respiratory Research Centre, The Kids Research Institute Australia, The University of Western Australia, Perth, Western Australia, Australia; UWA Centre for Child Health Research, The University of Western Australia, Perth, Western Australia, Australia; The Institute For Respiratory Health, The University of Western Australia, Perth, Western Australia, Australia; Occupation, Environment and Safety, School of Population Health, Curtin University, Perth, Western Australia, Australia; School of Biomedical Sciences, The University of Western Australia, Perth, Western Australia, Australia, Australia; Department of Respiratory and Sleep Medicine, Perth Children’s Hospital, Perth, Western Australia, Australia; Centre for Cell Therapy and Regenerative Medicine, School of Medicine and Pharmacology, The University of Western Australia and Harry Perkins Institute of Medical Research, Perth, Western Australia, Australia

## Abstract

*Pseudomonas aeruginosa* is a priority pathogen in chronic and multidrug-resistant infections, yet therapeutic phages targeting this organism often exhibit variable and unpredictable efficacy. A mechanistic understanding of the genomic determinants governing phage–host interactions is therefore critical for the rational design of robust phage therapeutics. Here, we systematically dissected the genetic featrures underlying lytic outcomes across a diverse panel of *P. aeruginosa*-infecting phages. We comprehensively annotated receptor-binding proteins (RBPs), bacterial defence systems and phage-encoded anti-defence genes, and experimentally defined host receptor usage for each phage, linking receptor specificity to cognate RBPs. We identified multiple anti-defence systems—including *vcrx089*, acrIIA24, *atd1*, *gnarl1*, *klcA*, *darA* and *nmna* (NARP2-associated)—and RBPs targeting lipopolysaccharide, type IV pili and flagella that are associated with enhanced lytic activity and represent tractable engineering targets.

Across 174 genomic features analysed, 110 significantly influenced phage activity, with heterogeneous and context-dependent effects. Leveraging these features, we trained a machine-learning classifier that accurately predicted phage–host outcomes (AUC–ROC = 0.875), demonstrating that interaction phenotypes are encoded in definable genomic signatures.

Together, our findings reveal the quantitative contribution of phage anti-defence systems to infectivity in *P. aeruginosa* and define a genomic framework for predicting and engineering lytic success. These results establish a foundation for the rational design of synthetic phages with enhanced host ranges.

## Introduction

The increase in antimicrobial-resistant (AMR) bacterial infections poses a growing threat to modern healthcare (O’Neill 2016). AMR is projected to cause up to 10 million deaths annually by 2050, and recent estimates already attribute 4.95 million deaths to drug-resistant infections in 2019, indicating that current trends are consistent with this trajectory (O’Neill 2016, Murray et al. 2022). Despite this growing burden, the antibiotic development pipeline remains insufficient, underscoring the urgent need for alternative antimicrobial strategies (Theuretzbacher et al. 2020, Reig et al. 2022, WHO 2022).

The use of bacteriophages (phages) is an increasingly promising alternative antimicrobial strategy in the post-antibiotic era (Aslam et al. 2020, Pirnay et al. 2024, Weiner et al. 2025). Although conceptually straightforward, phage infection involves a complex interplay between phage and bacterial factors (Georjon et al. 2023, Burke et al. 2024, Costa et al. 2024, Gaborieau et al. 2024, Müller et al. 2024, Murtazalieva et al. 2024). Infection begins at the bacterial surface, where phages must recognise and bind specific receptors via receptor-binding proteins (RBPs; Burke et al. 2024, Gaborieau et al. 2024, Müller et al. 2024). Following adsorption, the phage injects its genome into the host cell, where infection may be blocked by bacterial defence systems, including degradation of phage DNA, abortive infection pathways, or interference with phage replication and assembly (Georjon et al. 2023). In response, phages may encode anti-defence mechanisms that counteract these systems through multiple strategies, such as shielding phage components from recognition, inhibiting defence protein activity or complex formation, sequestering or degrading defence signalling molecules, or reconstituting phage components that are actively degraded by the host (Murtazalieva et al. 2024). However, natural phages are constrained by the limited repertoire of anti-defence mechanisms they can encode (Costa et al. 2025). This ongoing molecular arms race often results in a narrow host range, which can be advantageous for precision-based therapy by minimising off-target effects on the microbiota (Mu et al. 2021, Pirnay et al. 2024, Weiner et al. 2025). Conversely, restricted host range presents practical challenges, as it limits the coverage of individual phages and necessitates time-consuming matching to patient isolates (Pirnay et al. 2024, Weiner et al. 2025). Genetic engineering of phages to incorporate factors termed “lytic determinants” represents a promising strategy to overcome these limitations and enable the development of next-generation phage therapeutics with expanded or modulated host ranges (Ando et al. 2015, Huss et al. 2021, Latka et al. 2021, Qin et al. 2022, Garb et al. 2025, Yamashita et al. 2025).

Here, we sought to identify key bacterial defence systems, phage-encoded anti-defence mechanisms, and RBPs that collectively determine phage susceptibility, using *Pseudomonas aeruginosa* as an exemplar organism to inform rational phage engineering. *P. aeruginosa* is among the most challenging AMR pathogens due to its high levels of intrinsic and acquired resistance and virulence, which contribute substantially to morbidity, mortality, and increasing healthcare burden (Murray et al. 2022, Wozniak et al. 2022, WHO 2024). Previous investigations of phage lytic determinants have produced conflicting conclusions regarding the contribution of *P. aeruginosa* defence systems to phage susceptibility and have largely overlooked the role of phage-encoded anti-defence mechanisms (Burke et al. 2024, Costa et al. 2024, Müller et al. 2024).

To address this gap, we assembled a diverse panel of 29 phages and 88 clinical *P. aeruginosa* isolates, annotated candidate lytic determinants, characterised the bacterial receptors targeted by RBPs, and quantified the contribution of each determinant to phage activity. This integrated analysis enabled the identification of RBPs, anti-defence factors, and host defence systems that are critical drivers of phage–*P. aeruginosa* interactions and therefore represent tractable targets for genetic engineering. We further validated these findings by using the identified key lytic determinants to train a machine learning classification model that accurately predicted phage activity against *P. aeruginosa* isolates *in silico*. This approach not only supports the predictive value of the identified genomic features but also provides a practical framework to guide phage selection and design. To our knowledge, this represents one of the most comprehensive analyses of lytic determinants in *P. aeruginosa*–phage systems, integrating empirical phenotypic data with predictive modelling. Collectively, these findings establish a foundation for the rational design of next-generation phage therapeutics with expanded host range and improved reliability for clinical application.

## Methods

Please refer to supplementary material for full details.

### Bacterial isolates

The laboratory *P. aeruginosa* strain PAO1 was kindly provided by Professor Barbara Chang (University of Western Australia, Australia), and isogenic mutants were obtained from the Manoil laboratory (University of Washington, USA). Clinical *P. aeruginosa* isolates were supplied by Dr Anna Tai (Sir Charles Gairdner Hospital, Western Australia, Australia), Professor Sarath Ranganathan and Ms Rosemary Carzino (Murdoch Children’s Hospital, Victoria, Australia), and Professor Geoffrey Coombs (Murdoch University, Western Australia, Australia). Isolates were received de-identified and stored at −80 °C in 25% (v/v) glycerol. Routine culture was performed at 37 °C on Luria–Bertani (LB) agar (Becton Dickinson, USA), with single colonies propagated overnight in LB broth with agitation (120–200 rpm).

### Bacterial DNA extraction, whole-genome sequencing and bioinformatic analyses

A total of 110 isolates were cultured overnight, and approximately 10^9^ colony-forming units were pelleted by centrifugation (4,000 × g, 10 min). Genomic DNA was extracted using the Nanobind CBB kit (PacBio Biosciences, USA) according to the manufacturer’s protocol and quantified using the Qubit dsDNA Broad Range assay (Thermo Fisher Scientific, USA), with integrity assessed by 1% agarose gel electrophoresis. Long-read sequencing was performed on the Oxford Nanopore PromethION 2 Solo platform (Oxford Nanopore Technologies, UK). Libraries were prepared using the Native Barcoding Kit 96 V14 (SQK-NBD114.96) and sequenced on FLO-PRO114M (R10.4.1) flow cells. Basecalling and demultiplexing were performed with Dorado (v0.5.1) using the dna_r10.4.1_e8.2_400bps_sup@v4.3.0 model, and BAM files were converted to FASTQ format with samtools (v1.43.0; Li et al. 2009). Read quality control was performed with Kraken (v1.1.1) using the MiniKraken database (Wood et al. 2014), followed by filtering with Filtlong (v0.2.1) and visualisation with NanoPlot (v1.42.0; De Coster et al. 2023). Assemblies were generated using Flye (v2.9.3), plasmids were predicted with rfplasmid (v0.0.18; van der Graaf-van Bloois et al. 2021), and targeted plasmid assembly was performed with plassembler (v1.6.2; Bouras et al. 2023). Chromosomal and plasmid contigs were merged and reordered using Dnaapler (v0.3.1; Bouras et al. 2024). Assembly quality was assessed with CheckM (v1.2.2; Parks et al. 2015), minimap2 (v2.28; Li 2018), samtools (Li et al. 2009), Qualimap (v2.3.0) and QUAST (v5.2.0; Mikheenko et al. 2015, Okonechnikov et al. 2016). Genomes were annotated using Bakta (v1.9.3; Schwengers et al. 2021). Specialised annotations included Multilocus sequence typing (MLST; mlst v2.23.0; Seemann 2022), prophage prediction (PhiSpy v4.2.21; Akhter et al. 2012) and defence system identification (PADLOC v2.0.0; Payne et al. 2021). Serotypes were assigned using PAst (Thrane et al. 2016).

Isolates were excluded on the basis of contamination, poor assembly quality, minimal SNP divergence (<10 SNPs; snp-dists; Supplementary Table 1) or consistently poor growth. Two closely related isolates (M1C010 and M1C124) were retained owing to them being the only in-house members of *Pseudomonas paraeruginosa*. The final dataset comprised 86 *P. aeruginosa* and two *P. paraeruginosa* isolates used for downstream analyses. For pangenomic analyses, reference genomes from Ozer *et al*. (2019) were downloaded using NCBI Datasets (O’Leary et al. 2024). The pangenome was inferred with PPanGGOLiN (v2.0.5), and phylogeny reconstructed from the pan genome alignment using IQ-TREE (v2.0.3; JTT+F+I+G4; 1,000 ultrafast bootstraps; Gautreau et al. 2020). Trees were visualised with iTOL (Letunic et al. 2021). Defence system subfamily count and presence–absence matrices were extracted from PADLOC outputs for downstream analyses and M1C124 and M1C010 were excluded as separate species.

### Bacteriophage isolation, visualisation and nomenclature

Phages were isolated from wastewater collected at the Subiaco Wastewater Treatment Plant (Shenton Park, Western Australia, Australia) as described previously (Ng et al. 2022, Vaitekenas et al. 2022). Phages were propagated and titrated using double-overlay agar assays. Virions were visualized by transmission electron microscopy at the University of Adelaide (Iszatt et al. 2025; Supplementary Fig. 1). Since isolates were obtained on Noongar Wadjuk Boodjar, candidate phages were assigned descriptive names following consultation with the Noongar Language Centre (Perth, Western Australia; Supplementary Table 2).

### Phage DNA extraction, sequencing and annotation

Phage DNA was extracted as previously described and sequenced by the Australian Genome Research Facility (Melbourne, Australia; Jakočiūnė et al. 2018). Libraries were prepared using the Nextera XT kit (Illumina) and sequenced on a NovaSeq 6000 platform (150 bp paired-end). Assemblies and phage contig identification were performed using the Phanta pipeline (v0.3; Iszatt 2023). Completeness and contamination were assessed using read-mapping statistics and CheckV (Nayfach et al. 2021).

Anti-defence systems were annotated using DefenseFinder (v1.0.10) and dbAPIs, with high-confidence calls retained at e-value ≤ 1 × 10^−5^ (Yan et al. 2023, Tesson et al. 2024). Where annotations differed, DefenseFinder assignments were prioritised. RBPs were predicted using PhageRBPdetection (v4), and annotations corresponding to anti-defence or lytic effector proteins were excluded (Boeckaerts et al. 2022). Presence–absence matrices were constructed for RBP and anti-defence repertoires.

### Generation and sequencing of phage-resistant mutants

A PAO1 mutant resistant to *Phikzvirus* J5TC was generated and purified as described previously (Wannasrichan et al. 2022). DNA was extracted using the DNeasy Blood and Tissue kit (Qiagen) and sequenced by the Australian Genome Research Facility (AGRF; Melbourne, Victoria, Australia). Variants relative to PAO1 were identified using Snippy (v4.6.0; Seeman 2020; Supplementary Table 3).

### Efficiency of plating and receptor determination

Putative receptors were inferred using modified efficiency-of-plating (EOP) assays (Khan Mirzaei et al. 2015). Serially diluted phages were spotted onto overlay agar containing isogenic PAO1 mutants or in-house derivatives, followed by incubation, titre determination and EOP calculation (Gibson et al. 2019).

### Minimum lytic concentration (MLC)

MLC was determined as previously described with minor modifications (Gaborieau et al. 2024). Phages were diluted to 10^8^ and 10^6^ PFU ml^−1^ and spotted onto lawns of each of the 88 isolates. Plates were incubated overnight at 37 °C and scored for lysis. MLC values were assigned according to the lowest concentration producing detectable lysis.

### Host-range structure analyses

Interaction matrices were binarised (lytic, MLC > 1). Pairwise Jaccard distances between phages were calculated, and associations with phage genus or isolation host were tested using PERMANOVA (999 permutations). Dispersion effects were assessed with PERMDISP. Analyses were performed in Python (scikit-bio v0.6.3; Jai Ram Rideout 2025).

### RBP diversity and host-range association

RBP richness and Shannon entropy were calculated from filtered RBP matrices. Associations with infectivity were tested using Spearman correlation. Pairwise RBP and host-range dissimilarities were compared using Mantel tests (999 permutations). Individual RBPs were ranked by correlation with infectivity.

### Defence systems and infectivity

Defence burden was related to infectivity using mixed-effects models with defence count as a fixed effect and phylogroup as a random intercept. Defence-specific effects were assessed using one-sided Wilcoxon rank-sum tests on isolate-level mean MLC values.

### Univariate and multivariate modelling

Genomic features were screened using Mann–Whitney U tests or Spearman correlations, with significant features (p < 0.05) retained. A CatBoost classifier was trained on filtered features using repeated stratified splits (85:15). Model performance was assessed using ROC–AUC, precision–recall curves, and confusion matrices with bootstrapped confidence intervals.

## Results

### A Panel of 88 Diverse *P. aeruginosa* Isolates that is Representative of the Recognised Population Structure

The in-house panel of clinical *P. aeruginosa* isolates (*n* = 88; Supplementary Table 4) captured diversity representative of globally circulating populations (Fig. 1). Comparison with previously defined population structures (Ozer *et al*., 2019) showed that the panel included representatives of the major clades A, B and C (Fig. 1a), which are broadly associated with the presence or absence of the exotoxins *ExoS* and *ExoU* (Supplementary Fig. 2). Clade assignment was not structured by serotype, continent of origin or clinical source (Supplementary Figs. 3–5), indicating that these epidemiological variables did not drive the observed phylogenetic relationships. Unsurprisingly, the two *P. paraeruginosa* isolates (M1C010 and M1C124) formed a distinct outgroup (Fig. 1a).

**Figure 1:**
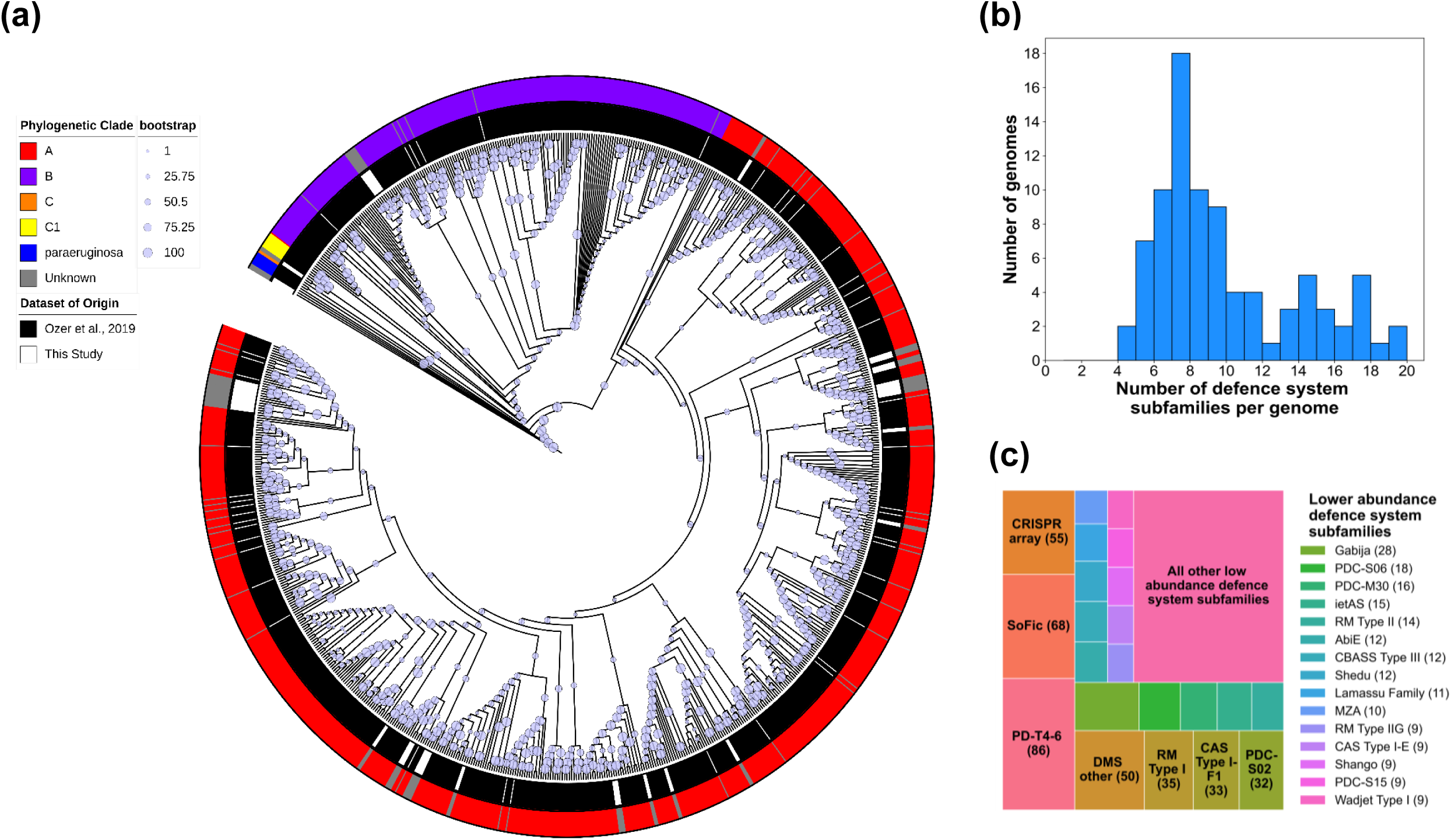
Characteristics of *P. aeruginosa* study isolates. **(a)** Pangenome phylogenetic tree of the *P. aeruginosa* isolates with 1,000 ultrafast bootstraps. Coloured rings indicate the phylogenetic clade (outer ring) and whether the isolates are from this study (white) or publicly available (Ozer et al., 2019, black; inner ring). The branch circle sizes correspond to bootstrap values as indicated in the legend. **(b)** Number of defence system subfamilies per bacterial genome **(c)** Frequency of bacterial defence systems subfamilies, where each square represents a system and the size corresponds to how many times it was identified. Numbers in parentheses reflect the number of isolates that contained each defence system. Colours indicate the system that was identified indicated in the legend.

To further characterise the diversity of the *P. aeruginosa* isolate panel, genomic features were comprehensively annotated. MLST identified 39 isolates belonging to previously unreported sequence types, whereas the remaining 47 isolates were assigned to 40 distinct published MLSTs (Supplementary Fig. 6). Among these, 36 of 40 sequence types were represented by single isolates, with only ST242, ST253, ST395 and ST591 observed more than once, indicating limited clonal expansion within the panel. Substantial diversity was also evident in lipopolysaccharide (LPS) serotypes, with 10 of the 20 recognised *P. aeruginosa* serotypes detected (Supplementary Fig. 7). The distribution was dominated by O6 (*n* = 33), followed by O1 (*n* = 15) and O3 (*n* = 15), whereas all remaining serotypes were infrequent (<10 isolates each).

Isolates additionally encoded a broad repertoire of defence systems, comprising 813 unique subfamilies, with a median of eight defence-system subfamilies per strain (range 4–19; Fig. 1b). Of these, 75 (9.2%) were classified as putative defence candidates (PDCs; Supplementary Table 4). The most prevalent systems included PD-T4-6 (present in all isolates), SoFic (68 isolates), CRISPR arrays (55 isolates) and DMS-other (50 isolates), whereas all other systems were detected in fewer than half of the isolates (Fig. 1c). Collectively, these data indicate that the isolate panel captures extensive phylogenetic, serotype and defence-system diversity, providing a genetically heterogeneous framework for interrogating determinants of phage susceptibility.

### A Panel of 29 diverse *P. aeruginosa* Phages

A total of 220 *P. aeruginosa* phages were isolated to establish an in-house reference library (Supplementary Table 5). Comparative genomic analysis against publicly available *Pseudomonas* phage genomes resolved 21 discrete clusters alongside a larger, more complex network structure (Fig. 2). The in-house isolates were distributed across 10 of the 21 clusters and were also represented within the broader network, indicating substantial genomic diversity within the collection (Fig. 2).

**Figure 2:**
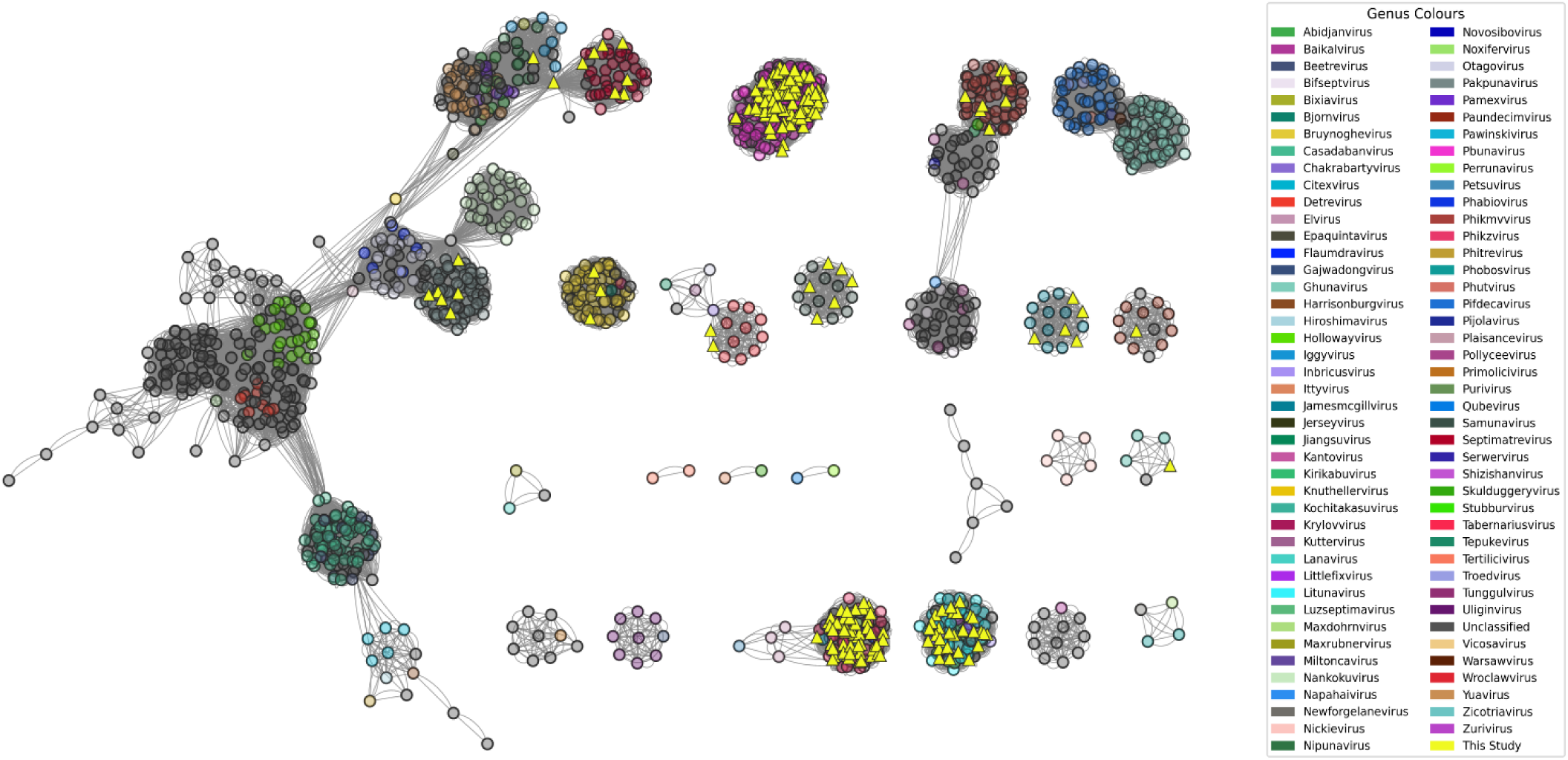
vConTACT2 network of shared genes among INPHARED (2^nd^ August 2024) *Pseudomonas* phages and our 251 in-house isolated *P. aeruginosa* phages. Node colours represent the genera indicated by the legend. Nodes are circular for publicly available phages and triangular for our isolated phages. Edges represent gene sharing between phage genome nodes.

A subset of 29 phages was selected for downstream characterisation to capture the phylogenetic and genomic breadth of the library. This panel included representatives from all recognised genera with the exception of *Jamesmcgillvirus*, which was excluded owing to the presence of multiple genera with comparable genome sizes (∼43 kb; Table 1). Where possible, phages spanning small, intermediate, and large genome sizes within each genus were included to maximise intrageneric diversity (Supplementary Table 5). Given the overrepresentation of *Pbunaviruses* and *Phikzviruses* within the collection, additional selections were made to prioritise genomic diversity within these groups (Table 1; Supplementary Table 5).

**Table 1:** General Genome Characteristics of the 29 Phages in the Diversity Panel.

| Phage | Length (bp) | GC% | CDS (n) | tRNA (n) | Putative Genus |
| --- | --- | --- | --- | --- | --- |
| EBID | 43,051 | 62.24 | 52 | 0 | *Phikmvirus* |
| 4TWA | 43,130 | 62.27 | 52 | 0 | *Phikmvirus* |
| WDPI | 43,175 | 54.54 | 54 | 1 | *Septimatrevirus* |
| YK3I | 45,460 | 52.62 | 70 | 4 | *Bruynoghevirus* |
| P71S | 49,457 | 44.75 | 68 | 0 | *Paundecimvirus* |
| V1IB | 64,200 | 60.33 | 89 | 0 | *Kochitakasuvirus* |
| N5HX | 62,547 | 55.59 | 81 | 0 | *Pbunavirus* |
| LG65 | 65,617 | 55.18 | 92 | 0 | *Pbunavirus* |
| 2CJA | 66,027 | 55.03 | 90 | 0 | *Pbunavirus* |
| C7E4 | 66,392 | 55.66 | 92 | 0 | *Pbunavirus* |
| 6281 | 67,446 | 55.81 | 93 | 0 | *Pbunavirus* |
| EUVX | 72,252 | 54.8 | 91 | 0 | *Litunavirus* |
| DKQ8 | 72,362 | 54.79 | 91 | 0 | *Litunavirus* |
| 0VBC | 73,338 | 54.67 | 94 | 0 | *Litunavirus* |
| KPSM | 89,617 | 49.27 | 162 | 15 | *Pakpunavirus* |
| 9XQE | 91,651 | 49.27 | 168 | 14 | *Pakpunavirus* |
| Z6TS | 92,794 | 49.4 | 168 | 15 | *Pakpunavirus* |
| AUV6 | 93,671 | 55.16 | 127 | 0 | *Samunavirus* |
| NC61 | 274,857 | 36.86 | 357 | 6 | *Phikzvirus* |
| R4QE | 277,163 | 36.81 | 357 | 8 | *Phikzvirus* |
| J5TC | 278,768 | 36.91 | 361 | 6 | *Phikzvirus* |
| NQ4L | 279,729 | 36.93 | 365 | 6 | *Phikzvirus* |
| T0U1 | 280,421 | 36.9 | 361 | 7 | *Phikzvirus* |
| MZOB | 281,074 | 36.85 | 358 | 4 | *Phikzvirus* |
| XMDS | 285,430 | 33.34 | 398 | 12 | *Wroclawvirus* |
| VKO7 | 287,890 | 33.22 | 405 | 12 | *Wroclawvirus* |
| GE9K | 302,312 | 43.61 | 386 | 8 | *Pawinskivirus* |
| O8XK | 305,647 | 43.62 | 386 | 8 | *Pawinskivirus* |
| 4NWX | 306,822 | 43.57 | 387 | 8 | *Pawinskivirus* |

### Bacteriophage Lytic Determinants

Receptor-binding proteins (RBPs) and anti-defence determinants influencing phage–host interactions were annotated across the phage panel (Supplementary Table 6). Sixteen anti-defence elements combined to create 16 distinct profiles were identified and grouped into nine functional systems: Anti-CRISPR/Cas (AcrII-24 and AcrVI-A1(Lwa)), Anti-TIR-STING (*atd1* and NTases), Anti-Gabija (*gad1*), the NAD reconstitution pathway (NARP1 and partial NARP2), O-antigen–associated barrier (*gnarl1*), anti-restriction–modification (*klcA*, *darA*, *vcrx089* and *adrA*), Anti-Retron (*Rad*), Anti-Thoeris (*tad1* and *tad1*- and *tad2*-like variants), and toxin–antitoxin (*mga47*; Fig. 3a). Most anti-defence loci were encoded as single genes, with the exception of NARP systems. Phages carrying NARP1 encoded both *adps* and *namat*, whereas those containing NARP2 encoded *nmna* with *nampt* absent (Fig. 3a).

**Figure 3:**
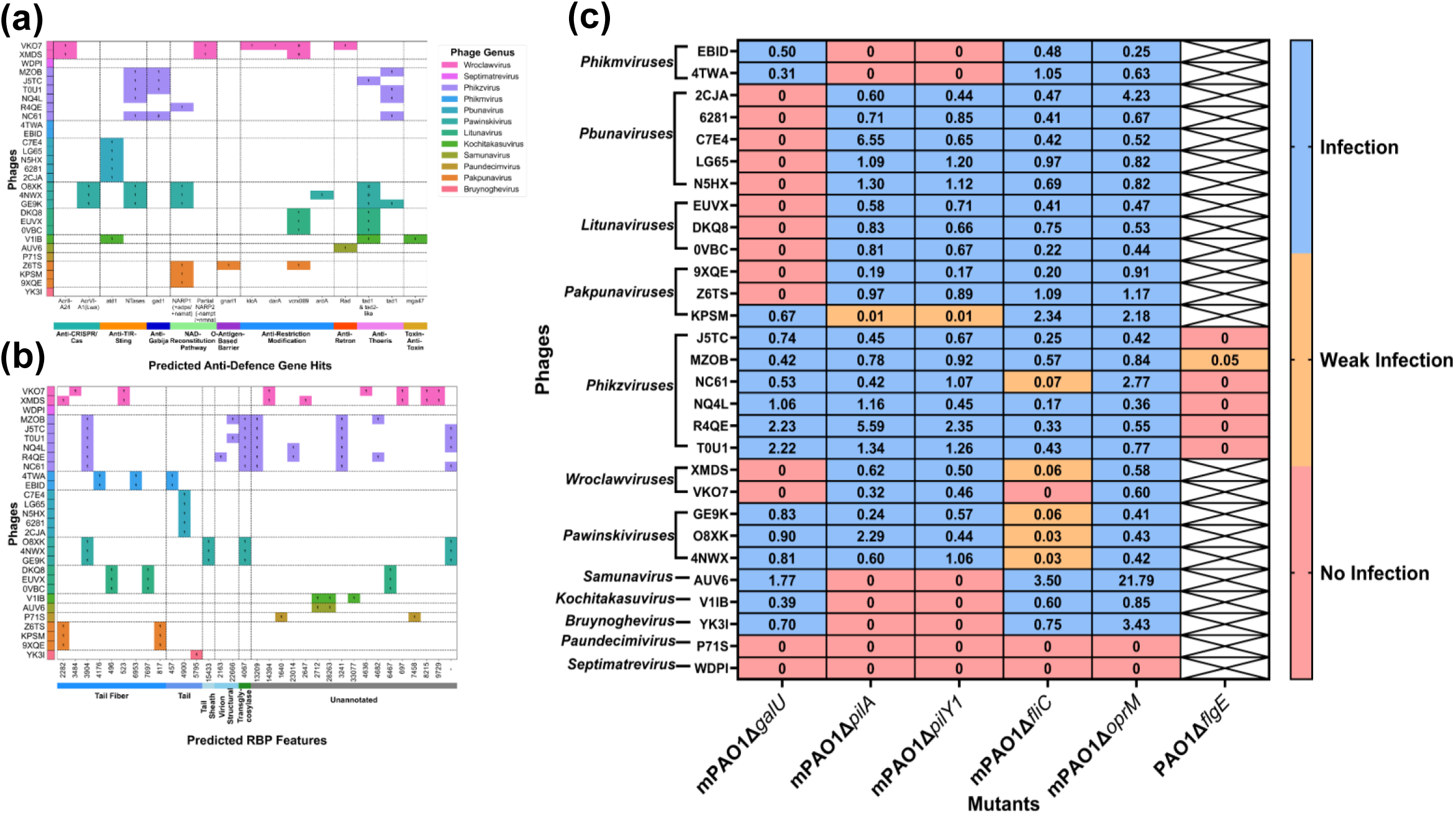
Empirical determination of phage receptor binding. **(a)** Heatmap of predicted anti-defences for the panel of 29 phages. Profiles are stratified by genus. Rows represent individual phages, coloured by their assigned genus (right), while columns represent predicted anti-defence genes grouped by functional category: Anti-CRISPR/cas (aqua), anti-TIR-sting (orange), anti-gabija (dark blue), NAD-reconstitution pathway (green), O-antigen-based barrier (purple), anti-restriction modification (light blue), anti-retron (red), anti-thoeris (violet) and toxin-anti-toxin (gold). **(b)** Heatmap of predicted receptor binding proteins (RBPs) for the panel of 29 phages. Rows are grouped by phage genus (indicated by colour blocks). Columns are numbered according to Protein Orthologous Group (PHROG) ID, and grouped by broad RBP type according to the following colour scheme: Structural (various hues of blue); host interaction (green), or unannotated (grey). In both figures **(a)** and **(b),** coloured cells denote gene presence, with colour reflecting phage genus as indicated in the legend. **(c)** Phage efficiency of plating (EOP) values for each of the mutants with altered surface structures. All EOP values were obtained by dividing the phage titre of the mutants by mPAO1 or PAO1. EOP was coloured according to no infection (EOP=0), weak infection (0<EOP<0.1) and infection (EOP≥0.1), denoted by red, orange and blue respectively.

Anti-defence genes were typically present as single copies within genomes; however, *gad1*, *tad1-*and *tad2*-like, and *vcrx089* occurred in multiple copies in a subset of phages (Fig. 3a). Anti-defence loci were sparsely distributed across the nine systems within the panel, although most genera encoded multiple systems. By contrast, *Pbunaviruses* and phage AUV6 each encoded only a single anti-defence, and *Phikmviruses*, WDPI, P71S and YK3I lacked high-confidence annotations (Fig. 3a). Intrageneric heterogeneity was evident, with only three of seven genera containing multiple phages with identical anti-defence profiles.

RBP repertoires were subsequently examined across the panel (Fig. 3b). RBP diversity was predominantly structured between genera, with intrageneric variation observed only in the *Wroclawvirus* and *Phikzvirus* genera. Individual phages encoded between 0 and 7 RBPs, grouped into 32 PHROG families, several of which remained unclassified. RBPs were typically present as single copies and were largely genus-specific, with limited intergeneric similarity, observed only between PHROG assignments in *Phikzviruses* and *Pawinskiviruses* and the *Samunavirus* and *Kochitakasuvirus* (Fig. 3b).

To assess whether variation in RBP composition corresponded to functional receptor usage, efficiency-of-plating (EOP) assays were performed using isogenic *P. aeruginosa* mutants (Fig. 3c). Phages from the *Pbunavirus* and *Litunavirus* genera were unable to infect the mPAO1Δ*galU* mutant (EOP = 0), indicating dependence on a correctly synthesised LPS core. Despite distinct RBP repertoires, *Pawinskiviruses* and *Wroclawviruses* exhibited markedly reduced or absent infection in the mPAO1Δ*fliC* mutant (EOP < 0.01 or 0), consistent with flagellotropic infection. *Wroclawviruses* additionally showed reduced infection in the mPAO1Δ*galU* mutant, suggesting that expanded RBP repertoires enable polyvalent receptor recognition.

Since *Phikzviruses* retained infectivity across the standard mPAO1 mutant panel, receptor mapping was extended using an in-house PAO1 derivative (Fig. 3c). Infection was abolished (EOP = 0) in a strain harbouring a large deletion in *flgE* (Supplementary Table 3), indicating flagellum-dependent infection. Four additional genera (*Phikmviruses, Bruynoghevirus, Kochitakasuvirus and Samunavirus*) required a functional type IV pilus, as evidenced by loss of infection in mPAO1Δ*pilA* and mPAO1Δ*pilY1* mutants. *Pakpunaviruses* utilised distinct receptors: phage, 9XQE which encode divergent RBP repertoires, showed weak infection in pilus mutants (EOP < 0.01), whereas phage Z6TS and 9XQE failed to infect mPAO1Δ*galU* (EOP = 0). Receptors for *Paundecimivirus* and *Septimatrevirus* could not be resolved owing to the absence of detectable infection across the panel. Together, these data indicate that the phage collection targets the major *P. aeruginosa* surface structures, including LPS, flagella and type IV pili, and demonstrate that phages encoding distinct RBP repertoires can converge on shared host receptors.

### Phage-Host Interaction Assessment

Phage infection outcomes were then determined through MLC and the investigation of 15,312 phage-*P. aeruginosa* interactions. Across three biological replicates for each phage-*P. aeruginosa* interaction, 70.43% showed complete consistency in the lysis scoring with 97.57% showing consistency in at least two of the three replicates. To contextualise the diversity of interactions within our *P. aeruginosa* panel, we first assessed the overall distribution of phage infectivity and bacterial susceptibility (Fig. 4a). Phages exhibited variable infectivity, lysing a median of 42 (range 1-69) of the 86 *P. aeruginosa* clinical isolates tested, and 41.4% of phages lysed more than half of the isolate panel (Fig. 4a).

**Figure 4:**
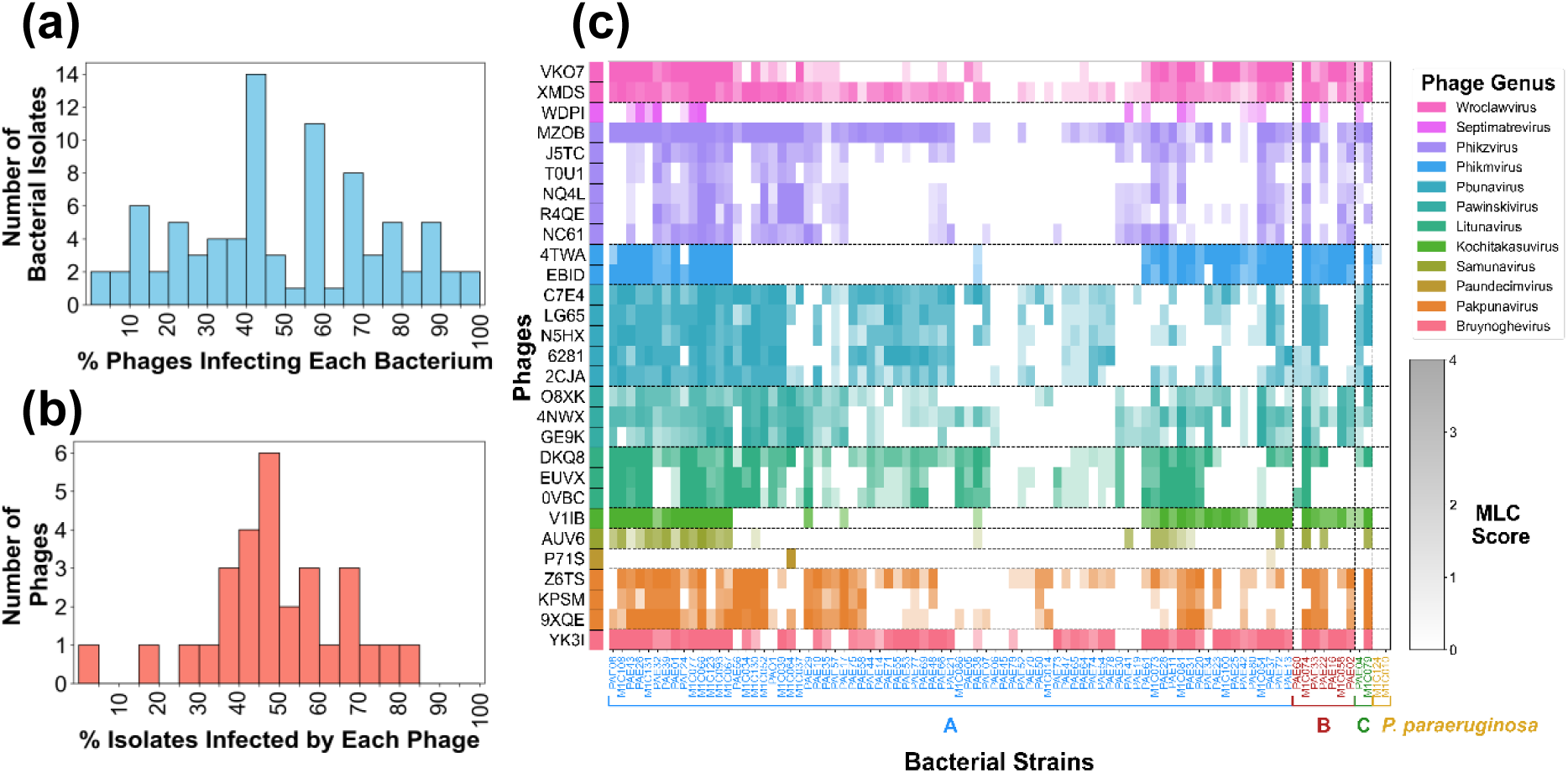
Phage and *P. aeruginosa*lytic interactions. **(a)** Histogram showing the number of bacterial isolates based on the percentage of phages capable of lysing them. **(b)** Histogram showing the number of phages based on the percentage of isolates they infected. **(c)** Heatmap of lytic activity (minimum lytic concentration, MLC) for 29 phages tested against 88 *P. aeruginosa* isolates. Rows are grouped by phage genus (indicated by colour blocks), and shading intensity within each row reflects MLC score using genus-specific colour gradients. Columns are grouped and coloured according to the established clades the *P. aeruginosa* isolates putatively belong to: A (blue), B (red), C (green), or outlier/unknown (gold).

Similarly, bacterial isolates displayed heterogeneous susceptibility profiles, with a median of 14 (range 0-28) phages capable of productive infection per isolate (Fig. 4b). The *P. aeruginosa* panel was quite permissive with 46.5% (40/86) of the isolates lysed by at least half of the phages (Fig. 4b). Despite the presence of broadly infective phages, WDPI (*Septimatrevirus*) and P71S (*Paundecimvirus*) exhibited highly restricted host ranges, lysing 15 and 1 isolates, respectively, well below the panel median host range of 43 (Fig. 4c). Similarly, while most *P. aeruginosa* isolates were permissive to at least one phage, PAE45 were not susceptible (MLC>1) to any of the 29 phages tested (Fig. 4c). Additionally the two *P. paraeruginosa* isolates (M1C010 and M1C124) were not susceptible to any of the phages tested (Fig. 4c).

### Multivariate Analyses of the Lytic Determinants Effects on Interaction Outcomes

Phage activity was first evaluated in the context of *P. aeruginosa* defence system content. The number of defence systems did not correlate with phage resistance (β_Num DS = −0.11, *p* > 0.05; Fig. 5a) or phage infections efficiency (β_Num DS = 0.00, *p* > 0.05; Fig. 5b). These data indicate that aggregate counts of defence systems does not measurably affect phage susceptible or efficiency of phages that successfully infect. Specific defence systems were next evaluated for evidence of broad resistance by ranking according to average MLC (Fig. 5c). None of the systems present in more than eight *P. aeruginosa* isolates significantly influenced mean MLC, suggesting that no individual defence system conferred broad-spectrum resistance across the panel.

**Figure 5.**
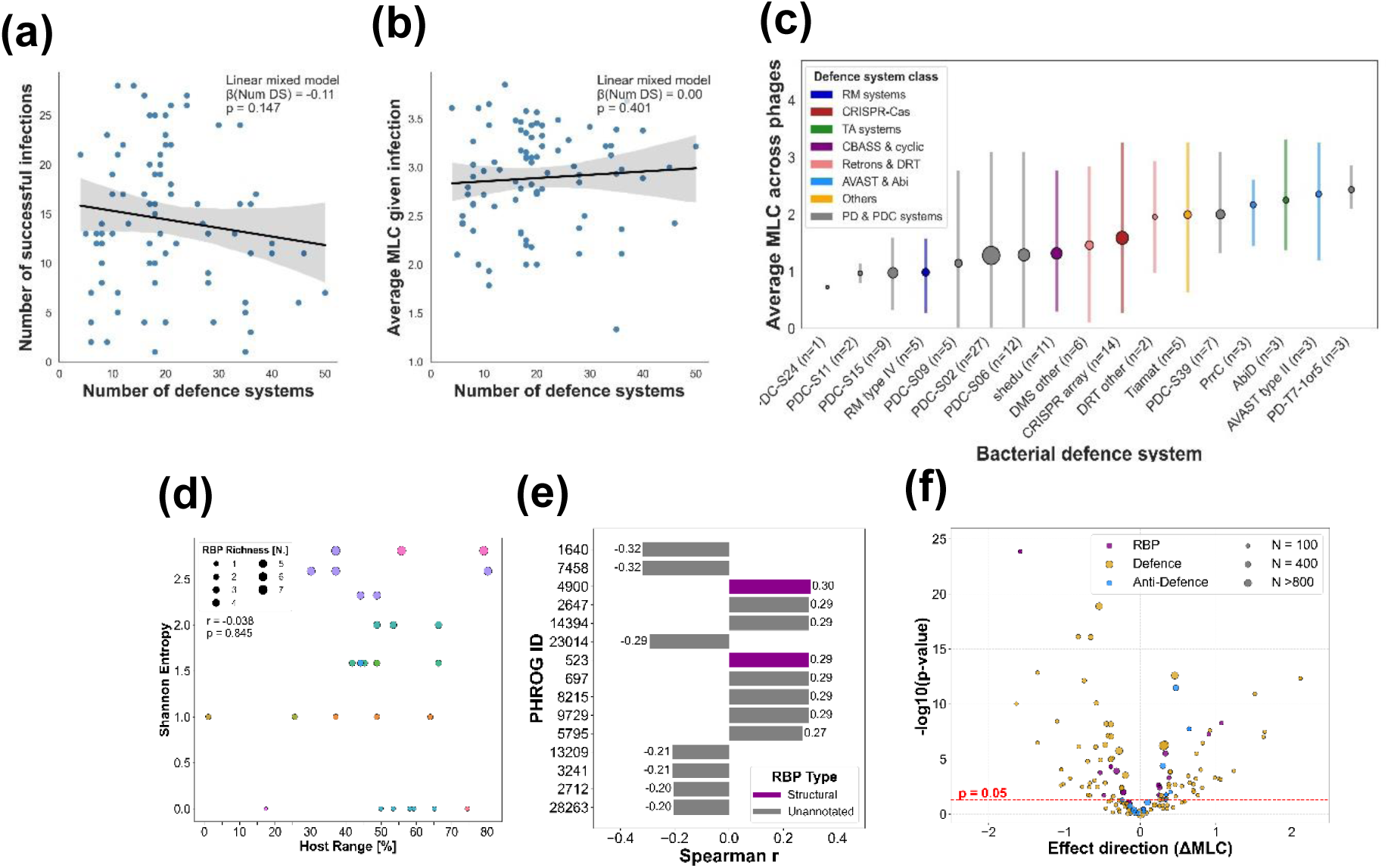
Influence of phage and *P. aeruginosa* lytic determinants on interactions. **(a)** Correlation of the number of defence systems per *P. aeruginosa* isolate with the number of phages capable of infecting them (MLC>1). **(b)** Correlation between the number of defence systems per *P. aeruginosa* isolate and its average MLC (only considers MLC>1). **(c)** Top 17 defence systems (x-axis) ranked by average MLC (y-axis). Defence systems are grouped by broad mechanistic category and coloured accordingly: RM systems – blue, CRISPR-Cas – red, TA systems – green, CBASS & cyclic – purple, Retrons and DRT – pink, HEC-like – gold, Zorya & Lamassu – teal, Others – orange, PD & PDC systems - grey. Mean is plotted as a circle proportional to the number of *P. aeruginosa* isolates the defence system was identified, along with the range. Systems present in >8 were tested for significant effect on average MLC via a Wilcoxon test. **(d)** RBP repertoire diversity correlation with phage host range. Each point represents a phage, coloured by genus using the same colour palette as previously defined in Figure 3a and scaled by host range (% of strains lysed). RBP Shannon entropy reflects diversity and evenness of those RBPs. **(e)** Top predicted RBPs correlated with phage infectivity. Spearman correlation coefficients (r) between the presence/absence of each predicted RBP and phage infectivity (defined as the percentage of *P. aeruginosa* isolates lysed). Bars are coloured by RBP type: structural (purple) or unannotated (grey). **(f)** Effect size versus statistical confidence for univariate predictors of phage infectivity. Each point represents a feature tested for its association with phage infectivity, as measured by the change in minimum lytic concentration (ΔMLC). The x-axis denotes the effect direction (mean difference in MLC between presence vs absence of the feature), while the y-axis shows statistical confidence (−log₁₀ p-value from Mann-Whitney U test). Marker colour indicates feature source: phage-encoded RBPs (purple), bacterial defence systems (gold), and anti-defence genes (blue). Marker size reflects the number of phage–bacteria combinations in which the feature was present. Only features with sufficient support (≥10 positive and ≥10 negative phage-*P. aeruginosa*) were considered.

Phage-intrinsic factors were subsequently examined to determine their contribution to activity patterns. Phage genus explained a substantial proportion of variation in host-range structure (pseudo-*F* = 7.06, *p* = 0.001), with an adjusted *R*² of 0.704 (Table 2). The bacterial host used for propagation also showed a significant but weaker association (pseudo-*F* = 1.91, *p* = 0.018; adjusted *R*² = 0.313). These effects were not attributable to differences in dispersion among groups (PERMDISP: genus *p* = 0.150; isolation host *p* = 0.950; Table 2). Together, these findings indicate that genomic features conserved within genera are the dominant determinants of host-range structure, whereas host-derived factors such as methylation exert a comparatively smaller effect.

**Table 2:** Summary of PERMANOVA and PERMDISP results assessing whether phage genus or isolation host explains differences in binary host range structure (presence/absence of lytic activity).

| Test | Grouping Variable | Sample Size | # Groups | Test Statistic | Adjusted $R^2$ | p-value | Method |
| --- | --- | --- | --- | --- | --- | --- | --- |
| PERMANOVA | Phage Genus | 29 | 12 | 7.06 | 0.704 | 0.001 | pseudo-F |
| PERMDISP | Phage Genus | 29 | 12 | 3.06 | - | 0.150 | F-value |
| PERMANOVA | Phage Host | 29 | 15 | 1.91 | 0.313 | 0.018 | pseudo-F |
| PERMDISP | Phage Host | 29 | 15 | 0.933 | - | 0.950 | F-value |

Given the strong genus-level structuring of infectivity and the conservation of receptor-binding proteins (RBPs) within genera, RBP repertoires were assessed as candidate determinants of host-range variation (Fig. 5d). No relationship was detected between overall RBP repertoire diversity (Shannon entropy) and infectivity breadth (Spearman *r* = -0.038, *p* > 0.05), with phages encoding larger RBP repertoires exhibiting infectivity profiles comparable to those with fewer RBP features.

Similarity in RBP composition was therefore evaluated as an alternative explanatory factor. Pairwise dissimilarities in RBP presence–absence profiles were compared with corresponding dissimilarities in binary host-range patterns. A Mantel test identified a weak but significant correlation between these matrices (Mantel *r* = 0.356, *p* = 0.001), indicating that phages with more similar RBP repertoires tend to exhibit more similar infectivity profiles. This relationship was supported by a complementary Spearman correlation analysis (Spearman *r* = 0.416, *p* < 0.001).

To explore contributions of individual RBPs, predicted RBP features were ranked according to their Spearman correlation with infectivity across the *P. aeruginosa* panel (Fig. 5e). Among the top 15 RBPs, eight showed positive associations with infectivity, consistent with a general contribution of RBPs to a broader host range. However, effect sizes were modest (*r* = −0.32 to +0.30), and the highest-ranking features were dominated by hypothetical proteins (13 of 15), with the remainder annotated as tail-associated proteins. Collectively, these analyses indicate that host-range structure is primarily shaped by genus-level genomic architecture, with RBP composition contributing to finer-scale variation in infectivity across *P. aeruginosa* isolates.

### Univariate Predictors of Phage Infectivity

To quantify the contribution of individual lytic determinants to phage infectivity, univariate association testing was performed across receptor-binding proteins (RBPs), bacterial defence systems, and phage anti-defence genes. For each feature, directional effects on infectivity were estimated as the change in mean MLC (ΔMLC), defined as the difference in mean MLC between phage–*P. aeruginosa* interactions in which the feature was present versus absent. Under this framework, ΔMLC > 0 indicated an association with increased phage activity, whereas ΔMLC < 0 indicated reduced activity (Fig. 5f).

Bacterial defence systems represented the largest class of features tested and exhibited a broad spectrum of effects on infectivity (Fig. 5f). In total, 78 defence systems were significantly associated with phage activity (*p* < 0.01), of which 42 were linked to reduced infectivity (ΔMLC < 0), consistent with their proposed roles in restricting phage infection. Notably, 36 defence systems showed significant positive associations with phage activity; however, these effects were generally modest, with only eight systems associated with effect sizes exceeding ΔMLC > 1 (*p* < 0.01).

A smaller number of RBPs ( *n* = 25) and anti-defence genes ( *n* = 7) were significantly associated with infectivity and typically displayed moderate effect sizes (ΔMLC between −1.0 and 1.0; Fig. 5f). Among RBPs, 16 of 25 showed positive associations with infectivity (*p* < 0.01), indicating enhanced activity when these features were present. The remaining nine RBPs were associated with reduced infectivity, although only two exhibited effect sizes below ΔMLC < −1 (upper left quadrant; Fig. 5f). Both RBPs originated from the *Paudecimivirus* P71S, which displayed highly restricted activity across the *P. aeruginosa* panel. All seven significant anti-defence genes (*vcrx089*, acrIIA24, *atd1*, *gnarl1*, *klcA*, *darA* and *nmna;* part of NARP2) were positively associated with infectivity (ΔMLC > 0; *p* < 0.01). Collectively, these analyses indicate that defence, anti-defence and receptor-binding determinants exert heterogeneous and gene-specific effects on phage–*P. aeruginosa* interactions, with bacterial defence systems contributing the largest share of variability in infectivity and phage-encoded features modulating activity within this constraint.

### Informed Classifier Modelling

To evaluate the combined predictive contribution of the identified lytic determinants (Fig. 5d), a CatBoost classifier was trained using these features to discriminate between lytic and non-lytic phage–host interactions. By modelling non-linear relationships and higher-order feature interactions, this framework enabled assessment of the integrative contribution of bacterial defence, receptor-binding and anti-defence determinants to infectivity outcomes.

The classifier demonstrated consistently strong performance, with a mean ROC–AUC of 0.875 (95% CI 0.846-0.906), indicating high discriminative capacity between lytic and non-lytic interactions (Fig. 6a). Precision remained high across moderate recall values, with an average precision of 0.87 (Fig. 6b). The distribution of ROC–AUC scores was tightly centred, supporting model stability and generalisability across validation folds (Fig. 6c). The mean confusion matrix indicated balanced classification performance, with 79% of lytic and 79% of non-lytic interactions correctly assigned (mean F1 score = 0.79; Fig. 6d).

**Figure 6.**
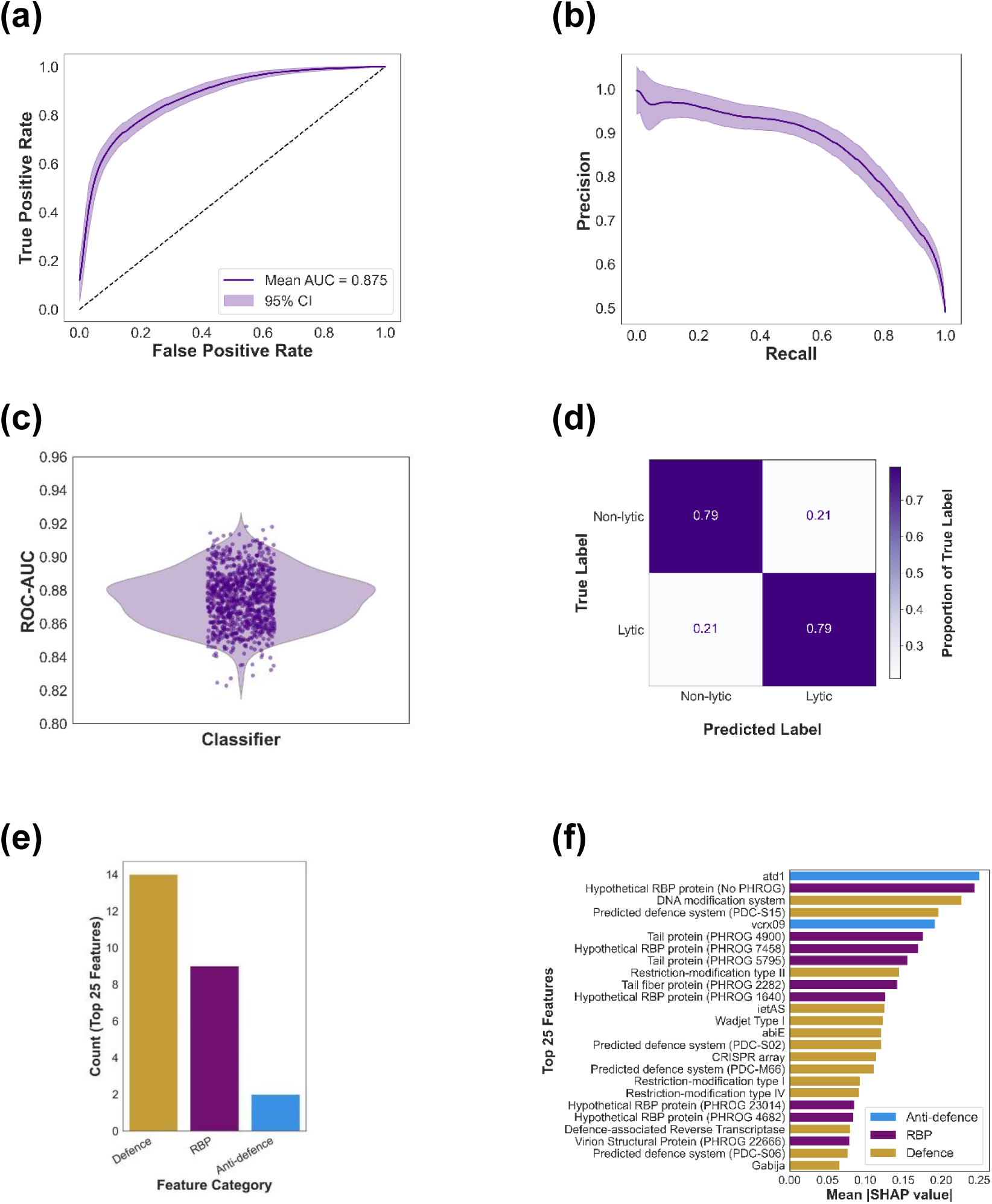
Performance of the CatBoost classifier in predicting phage lytic activity and top molecular predictors of phage infectivity. (a) Receiver Operating Characteristic (ROC) curve aggregated across 1000 stratified bootstrap iterations. The mean ROC curve is shown in purple with shaded 95% confidence intervals (mean AUC = 0.880, 95% CI: 0.845–0.913). (b) Precision–recall curve aggregated across the same bootstrap iterations. (c) Distribution of ROC-AUC values across bootstrap replicates. (d) Normalised confusion matrix averaged over bootstrap samples, illustrating the proportion of correctly and incorrectly classified lytic and non-lytic interactions. (e) Bar chart summarising the distribution of these top-ranked features by category. (f) Ranked summary plot of the top 25 features based on mean absolute SHAP values, reflecting their contribution to model predictions. Features are coloured by their assigned category: receptor-binding proteins (RBPs, purple), bacterial defence systems (gold), and anti-defence systems (blue).

To interpret model behaviour, SHAP values were computed and features ranked according to their mean absolute contribution to classification. Among the top 25 predictors, bacterial defence systems comprised the largest proportion (14 of 25), followed by RBPs (9 of 25) and anti-defence genes (2 of 25; Fig. 6e). The highest-impact features (SHAP > 0.15) included the anti-defence genes *atd1* and v*crx089*, PDC-S15 and DNA modification defence systems, restriction-modification system type II and several RBPs (PHROG 4900, 5795, 7458 and one unclassified; Fig. 6f). Together, these results indicate that variation in infectivity is best explained by the combined effects of bacterial defence architecture and phage-encoded receptor-binding and anti-defence modules, with defence systems contributing the dominant predictive signal.

## Discussion

These findings indicate that intracellular and extracellular determinants encoded by both phages and *P. aeruginosa* jointly shape interaction outcomes at a gene-resolved level. By comprehensively annotating lytic genetic determinants across a diverse panel of phages and *P. aeruginosa* isolates and linking these features to >15,000 phage–host combinations, multiple bacterial and phage genomic factors with variable effects on phage activity were identified. Notably, several phage-encoded determinants—including anti-defence genes (for example, *vcrx089*, acrIIA24, *atd1*, *gnarl1*, *klcA*, *darA* and the *nmna* gene of the NARP2 system) and receptor-binding proteins (RBPs) targeting the type IV pilus, lipopolysaccharide (LPS) core and flagellar structures—represent potentially actionable modules for engineering phages with expanded host range. The combined explanatory power of these determinants was supported by a machine-learning classifier capable of accurately discriminating lytic from non-lytic phage–host pairs (mean ROC–AUC = 0.875), indicating that the selected genomic features capture key biological signals underlying interaction outcomes.

Consistent with previous work, receptor binding emerged as a major determinant of phage activity (Müller et al. 2024). Here, this relationship was demonstrated across phages targeting multiple *P. aeruginosa* surface receptors, including LPS, type IV pili and flagella, suggesting that the association is generalisable across receptors. Sixteen RBPs were significantly associated with increased infectivity, including those representing the core RBP repertoires of *Pbunaviruses*, *Wroclawviruses*, *Litunaviruses* and the *Bruynoghevirus* included in this study. These repertoires collectively target the principal receptors exploited by *P. aeruginosa* phages and therefore represent candidate modules for host-range engineering, as previously proposed for synthetic receptor swapping approaches (Le et al. 2013, Ando et al. 2015, Latka et al. 2021). Conversely, nine RBPs were associated with reduced activity, including components of *Phikzvirus*, *Pawinskivirus*, *Samunavirus* and *Kochitakasuvirus* repertoires. These features may reflect structural constraints or ecological specialisation, although confirmation will require structural and adsorption studies (Le et al. 2013, Ando et al. 2015). The strongest negative associations corresponded to the *Paundecimivirus* RBP profile, consistent with the highly restricted host range of this genus.

The variable and occasionally counterintuitive effects of RBPs likely reflect strong environmental modulation of receptor interaction (Schumann et al. 2022, Ashworth et al. 2024, Li et al. 2025). Phage adsorption is known to depend on *P. aeruginosa* surface architecture, which is influenced by oxygen availability, nutrient status, and growth phase (Schumann et al. 2022, Ashworth et al. 2024, Li et al. 2025). RBPs negatively associated with activity under the present assay conditions may therefore confer advantages in alternative ecological contexts, whereas positively associated RBPs may be optimised for laboratory growth conditions. Future work incorporating adsorption assays across physiologically relevant environments will be required to resolve these context-dependent effects. Notably, overall RBP repertoire diversity was not associated with a broader host range. This contrasts with observations in Enterobacteriaceae phages, where receptor diversity correlates with expanded host breadth, and may reflect the narrower species specificity typical of *P. aeruginosa* phages (Sørensen et al. 2024).

In parallel, intracellular anti-defence mechanisms also contributed to variation in activity. Seven anti-defence genes were significantly associated with enhanced infectivity, suggesting that counter-defence modules can measurably influence interaction outcomes. Earlier studies reported limited effects of anti-defences on activity; however, these analyses were restricted by smaller datasets and narrower annotation scope (Costa et al. 2024). The present results suggest that anti-defence contributions become more apparent when diverse clinical isolates and broader defence repertoires are considered. Together, these findings indicate that extracellular receptor engagement and intracellular defence evasion operate as complementary determinants of phage success.

The influence of bacterial defence architecture was similarly context dependent. Previous studies have reported inconsistent relationships between defence system burden and phage susceptibility (Burke et al. 2024, Costa et al. 2024, Müller et al. 2024). Using a diverse clinical isolate panel, we did not detect an association between defence-system number and resistance frequency or phage infection efficiency, contrasting with earlier reports (Burke et al. 2024, Costa et al. 2024). Importantly, gene-level analyses revealed heterogeneous effects among individual defence systems, indicating that defence repertoires cannot be treated as functionally equivalent, nuances that aggregate analyses may miss (Burke et al. 2024, Costa et al. 2024). This variability is consistent with models in which many systems require cooperative activation or primarily target other mobile genetic elements (Marraffini et al. 2008, van Belkum et al. 2015, Bravo et al. 2024, Costa et al. 2024). These observations have practical implications for phage engineering. The combined abundance and impact of individual defence systems can be used to define the minimal counter-defence repertoire required for broad phage infectivity, which could be achieved through incorporation of natural or synthetic anti-defence modules (Qin et al. 2022, Garb et al. 2025, Yamashita et al. 2025).

Across the full dataset, 110 of 174 annotated lytic determinants significantly influenced phage activity and 94 features after a 85% training split were used to train a CatBoost classifier. The model demonstrated strong performance (ROC–AUC = 0.875; average precision = 0.87), comparable to previous predictive frameworks, but with improved precision. In contrast to studies in *Escherichia coli* that emphasised adsorption features, the present results indicate that intracellular defence and anti-defence mechanisms contribute substantially to predictive performance in *P. aeruginosa* (Gaborieau et al. 2024). This suggests that the relative importance of lytic determinants is host dependent, and that predictive models may require species-specific feature sets.

Several limitations warrant consideration. Interaction outcomes were measured under defined laboratory conditions and may not capture the full environmental variability encountered in clinical or ecological settings (Høyland-Kroghsbo et al. 2017, Høyland-Kroghsbo et al. 2018, Chen et al. 2025). In addition, the genomic framework does not account for gene expression dynamics, which are known to influence defence activation and phage replication (Høyland-Kroghsbo et al. 2018, Aframian et al. 2025, Li et al. 2025). Future studies incorporating transcriptomic or multimodal datasets may improve predictive resolution (Lood et al. 2022). External validation using independent *P. aeruginosa* isolate collections will also be necessary to assess generalisability.

In summary, specific anti-defence genes (*gnarl1*, *klcA*, *darA*, AcrIIA24, *nmna*, *atd1* and *vcrx089)* and RBP modules targeting the type IV pilus, LPS and flagella emerge as actionable determinants for engineering phages with expanded host range. A defined set of phage and bacterial genomic features was sufficient to train a robust classifier capable of predicting interaction outcomes in silico, supporting the feasibility of genome-informed phage matching. With further validation, this framework could facilitate rapid selection of candidate phages for personalised therapy and enable distributed, sequence-based matching of clinical isolates to globally available phage collections. More broadly, these findings illustrate how integrative genomic analysis, functional phenotyping, and machine learning can refine understanding of phage–host ecology while informing next-generation therapeutic design.

## Data Availability

All genomic data is available from NCBI and the accessions can be found in Supplementary Table 7. All other data is available upon request.

## Supporting information

Please refer to supplementary material for full details.

## Acknowledgements

We would like to acknowledge that this project was conducted and phages isolated on the traditional homelands of the Wadjak Noongar people. PhageWA thanks Sharon Gregory and Walyalap Waangkan Noongar language team, who named the phages in this study in Wadjak Noongar language. We would also like to acknowledge past and present PhageWA team members who helped isolate phages that have been analysed in this study. This includes Mr Scott Winslow, Ms Samantha Abigail McLean, Ms Jessica Hillas, Dr Daniel Rodolfo Laucirica and Dr Matthew Wee Peng Poh.

## Funding

This study was supported by a Future Health Research and Innovation Fund – Generative Artificial Intelligence Grant (IC2023-GAIA/21), an MRFF Chronic Respiratory Grant (2023559), as well as funding from the Stan Perron Charitable Foundation, the Conquer CF Foundation, and a Department of Health WACRF Grant. RNN is a BrightSpark Fellow and YK holds a CFWA Fellowship. S.M.S. is an NHMRC Practitioner Fellow holding an NHMRC Investigator Grant (2007725) and A.K. is a CFWA & Stan Perron Fellow.

## Declaration

None to declare.

## Author Contributions

AV (Conceptualisation, Investigation, Analysis, Writing-original draft). CJM (Investigation, Analysis, Writing-original draft, Writing-review and editing). LM (Investigation, Writing-original draft, Writing-review and editing). PGC (Analysis, Writing-original draft, Writing-review and editing). JJI (Conceptualisation, Analysis, Writing-review and editing). RNN (Writing-review and editing). STM (Investigation, Analysis, Writing-original draft, Writing-review and editing). YK (Analysis, Writing-review and editing). SMS (Writing-review and editing). AK (Funding Acquisition, Supervision, Conceptualisation, Analysis, Writing-review and editing).

## Notes

### Competing Interest Statement

The authors have declared no competing interest.

### Summary of Updates

The abstract has been revised to italicise species, genus and gene names. Author affiliations have been updated to include some that were missing.

