## Supplementary Fig. 1 for "Genomic Determinants of Phage Activity Against *Pseudomonas aeruginosa*: Roles of Receptors, Defence Systems, and Anti-Defences"

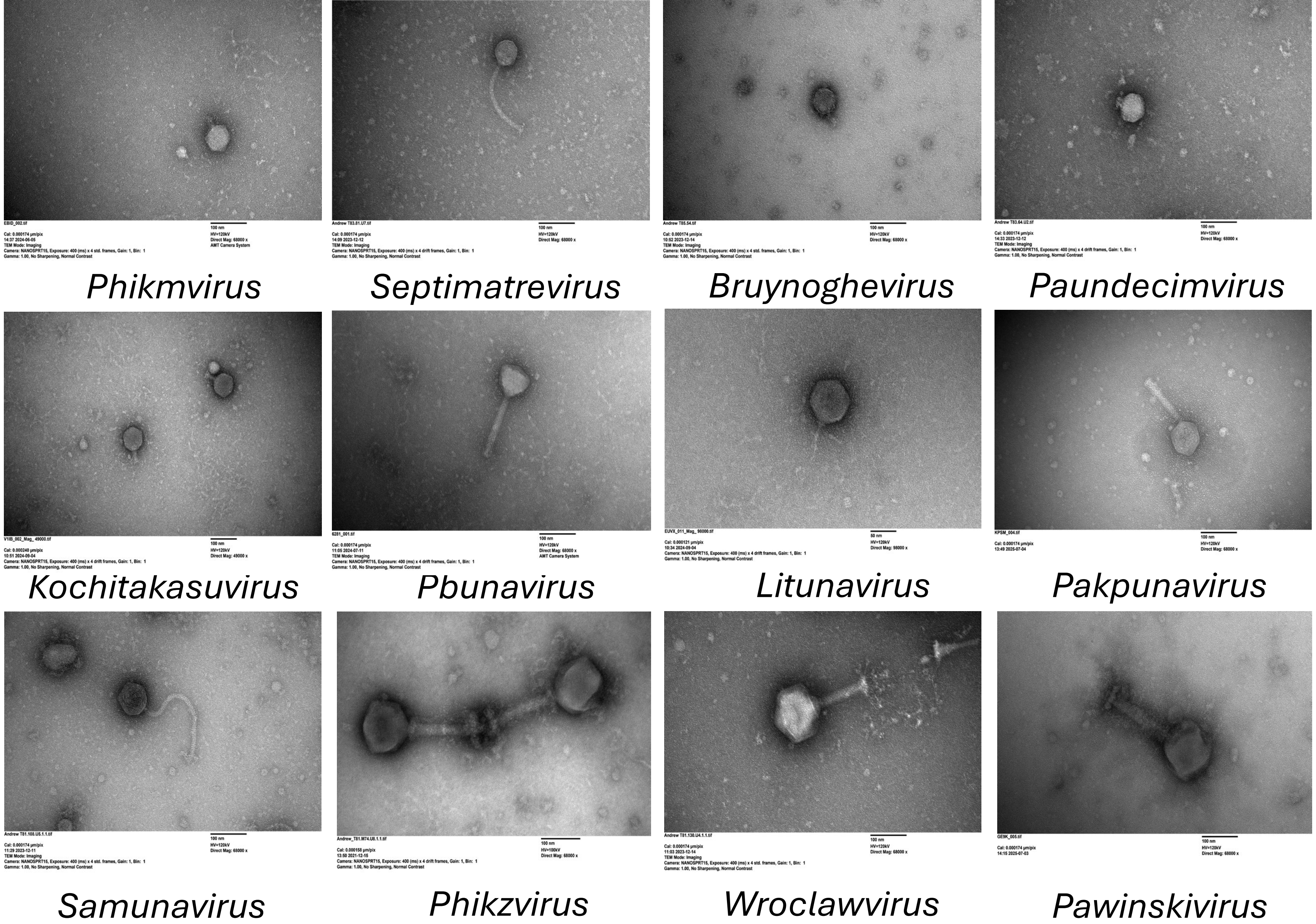


Supplementary Figure 1: Representative transmission electron microscopy (TEM) images of each phage genus that were used to give them descriptive names in the language of the Wadjak Noongar people.
