## Supplementary Figs. 3-5 for "Genomic Determinants of Phage Activity Against *Pseudomonas aeruginosa*: Roles of Receptors, Defence Systems, and Anti-Defences"

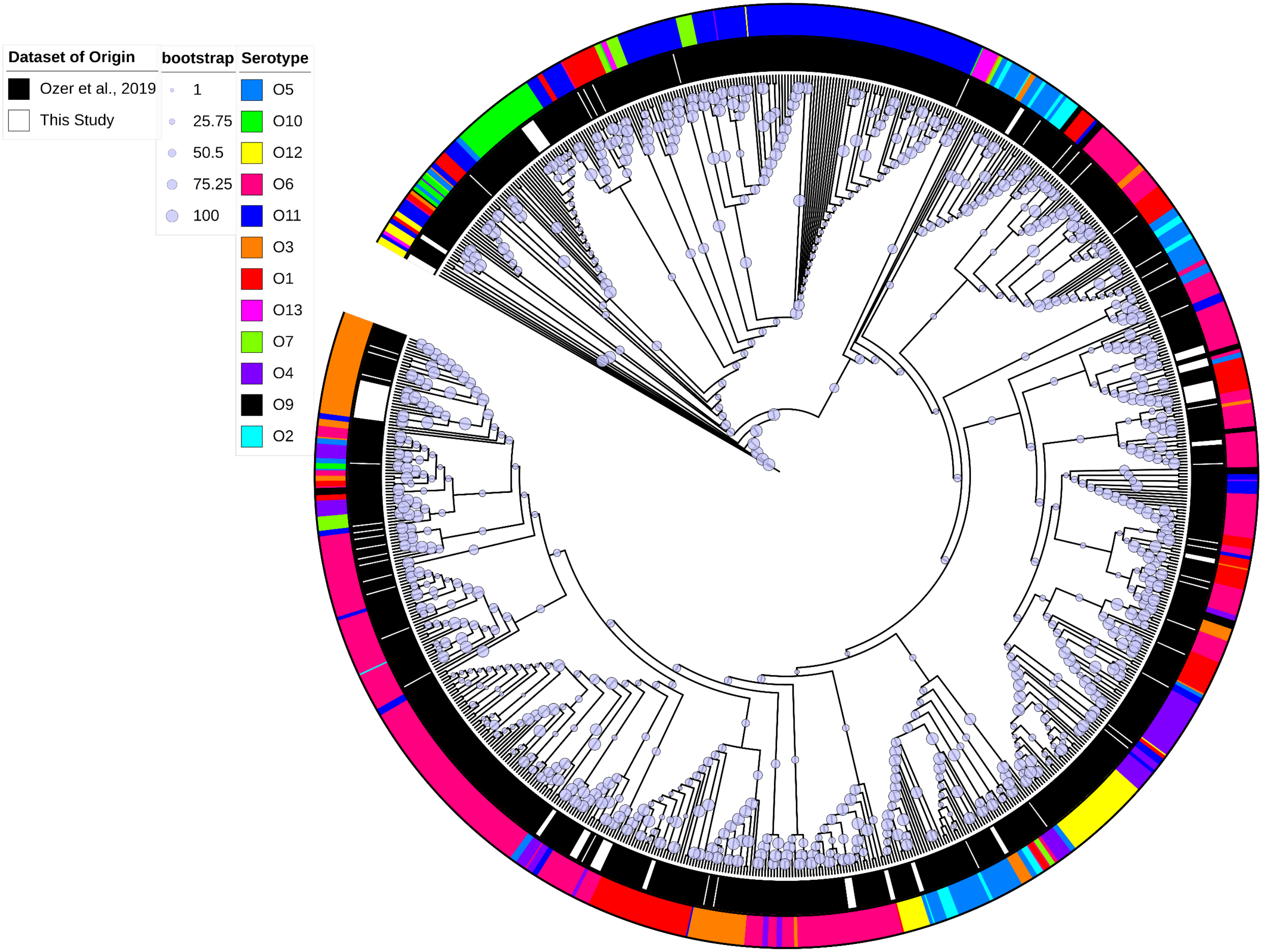


Supplementary Figure 3: Core Genome Phylogenetic Tree of the *P. aeruginosa* Isolates with 1,000 Ultrafast Bootstraps. The inner ring indicates if the isolates are from this study (white) or from Ozer et al., 2019 (black). The coloured outer ring indicates which serotype the isolates encode as depicted by the colours in the legend. The branch circle sizes correspond to bootstrap values as indicated in the legend.
