## Supplementary Fig. 6 for "Genomic Determinants of Phage Activity Against *Pseudomonas aeruginosa*: Roles of Receptors, Defence Systems, and Anti-Defences"

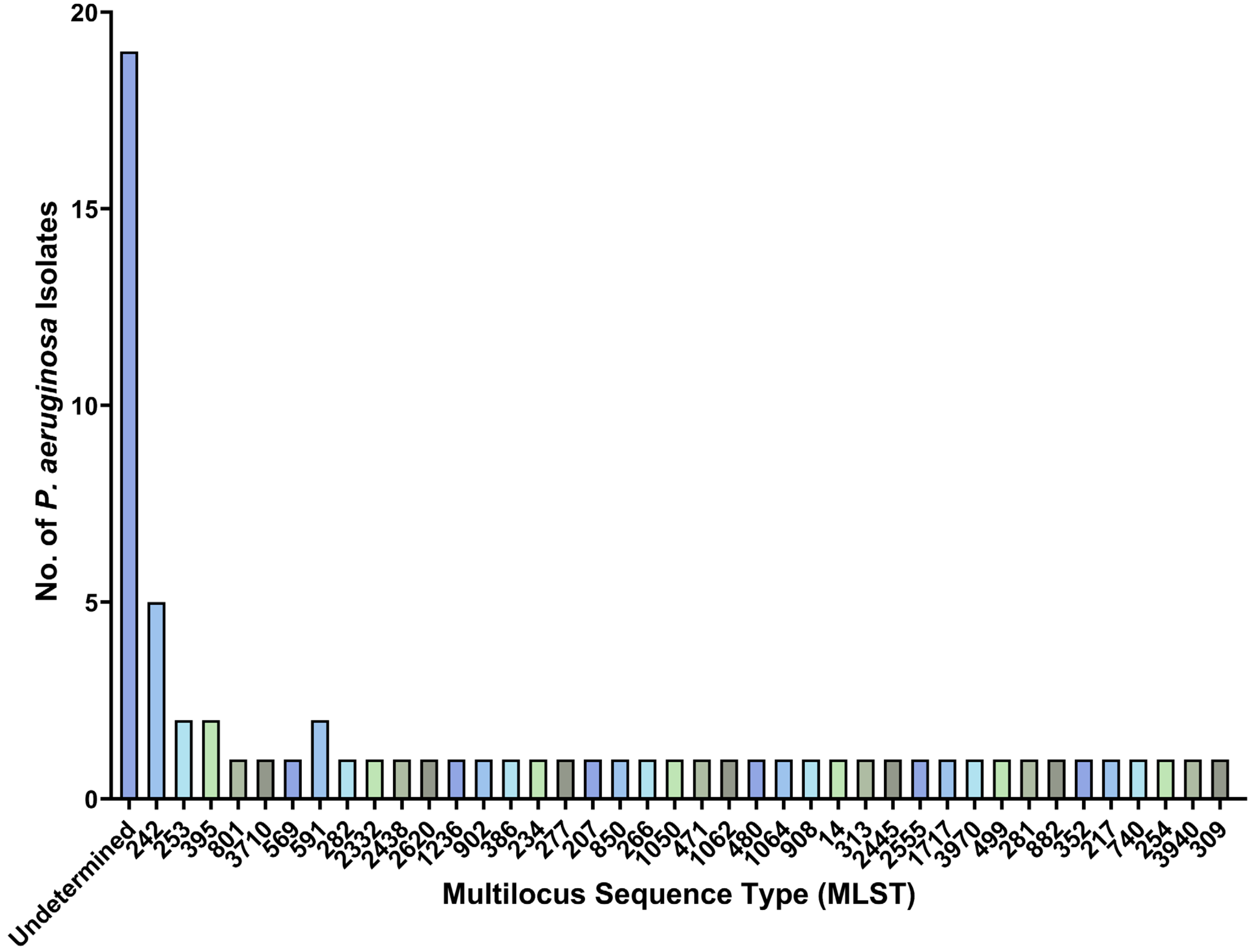


Supplementary Figure 6: The frequency of multilocus sequence types (MLST) of the *P. aeruginosa* in the 86 *P. aeruginosa* Isolate Panel.
