## Supplementary Fig. 7 for "Genomic Determinants of Phage Activity Against *Pseudomonas aeruginosa*: Roles of Receptors, Defence Systems, and Anti-Defences"

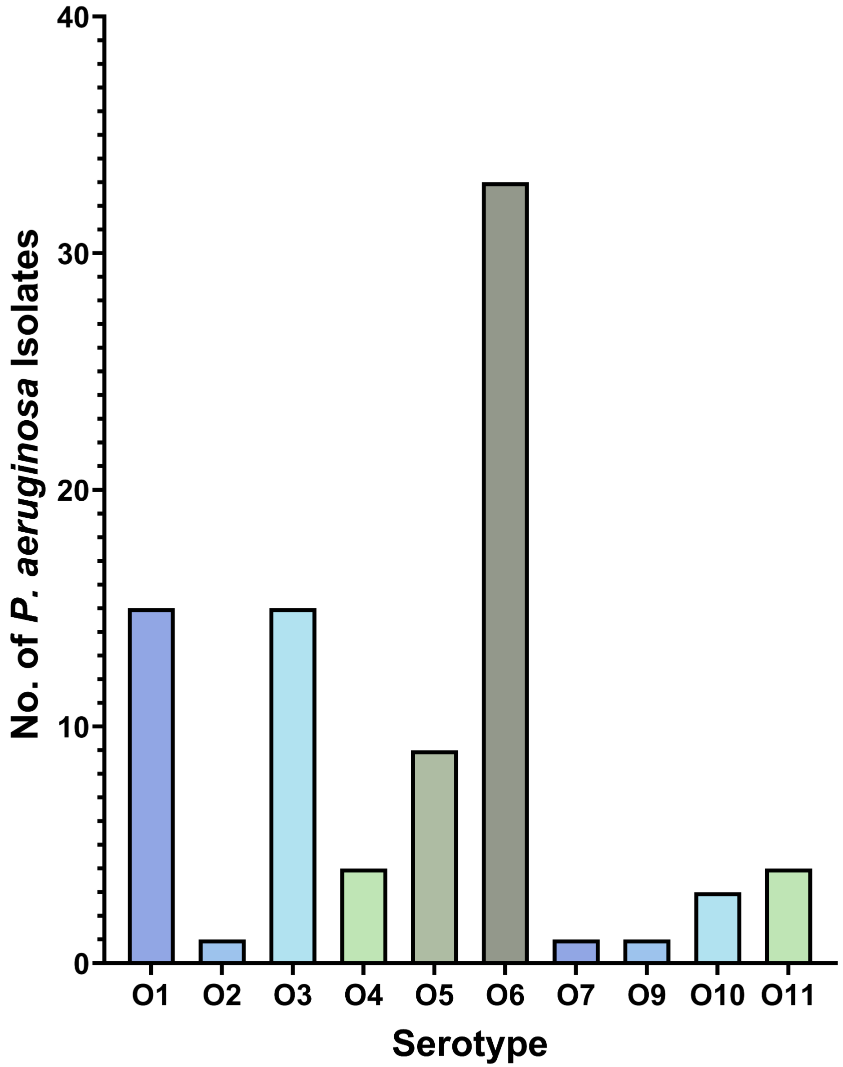


Supplementary Figure 7: The frequency of serotypes encoded by the *P. aeruginosa* in the 86 *P. aeruginosa* Isolate Panel.
