## Supplementary material for "Genomic Determinants of Phage Activity Against *Pseudomonas aeruginosa*: Roles of Receptors, Defence Systems, and Anti-Defences": Please refer to supplementary material for full details.

### Bacterial isolates

The laboratory *P. aeruginosa* strain, PAO1, was provided by Professor Barbara Chang (University of Western Australia, Australia) and isogenic mutants obtained from the Manoil laboratory (University of Washington, USA). Clinical *P. aeruginosa* bacterial isolates were provided by Dr Anna Tai (Sir Charles Gairdner Hospital, Western Australia, Australia), Professor Sarath Ranganathan and Ms Rosemary Carzino (Murdoch Childrens Hospital, Victoria, Australia) and Professor Geoffery Coombs (PathWest, Western Australia, Australia). Isolates were received, and frozen in 25% (v/v) glycerol at -80°C. *P. aeruginosa* was routinely cultured at 37°C on Luria-Bertani (LB) agar (Becton Dickinson, USA) and single colonies grown overnight in LB broth with 120-200 rpm agitation.

### Bacterial DNA extraction, whole genome sequencing (WGS) and bioinformatic analyses

Initially, 110 bacterial isolates were grown overnight and approximately $10^9$ colony forming units (CFUs) were pelleted by centrifugation for 10 minutes at 4,000 x *g*. Supernatant was discarded and DNA extracted from the pellet using the Nanobind CBB kit (PacBio Biosciences, USA) following manufacturer's instructions. DNA was quality controlled via Qubit dsDNA broad range assay (Thermo Fisher Scientific, USA) and 1% agarose gel electrophoresis.

Long-read WGS of bacterial DNA was performed using the Oxford Nanopore PromethION 2 Solo Oxford Nanopore Technolgies, UK). Libraries were prepared using the Native Barcoding Kit 96 V14 (SQK-NBD114.96; Oxford Nanopore Technologies, UK) and sequenced using PromethION 10.4.1 flow cells (FLO-PRO114M). Raw reads were basecalled and demultiplexed using Dorado (v0.5.1; Oxford Nanopore Technologies, UK) using the dna_r10.4.1_e8.2_400bps_sup@v4.3.0 basecalling model. Basecalled bam files were then converted to fastq files for downstream analyses using samtools v1.43.0 (1).

Quality control of reads was performed with Kraken v1.1.1 using the minikraken database created on 13/10/2017 (2). Reads were then filtered with filtlong v0.2.1 and visualised with NanoPlot v1.42.0 (3). Quality controlled reads were then assembled with Flye v2.9.3, and plasmids predicted with rfplasmid v0.0.18 (4). Targeted assembly of plasmids from quality-controlled reads was then performed with plassembler v1.6.2 and its database v1.5.0 (5). Chromosomal and plasmid contigs were then combined and reordered via Dnaapler v0.3.1 (6). Assemblies were assessed via Kraken v1.1.1 as described, checkM v1.2.2 and the 16/01/2015 database (7), minimap2 v2.28.0 (8), samtools v1.43.0 (1), qualimap v2.3.0 and Quast v5.2.0 (9, 10). The assemblies were then annotated with Bakta v1.9.3 with the full database v5.1.0 (11). Specialist annotation was performed with mlst 2.23.0 (12), Phispy v4.2.21 (13) and PADLOC v2.0.0 with its version 2.0.0 database (14). Annotation of each isolate's serotype was performed separately using the PAst script (15). X amount of *P. aeruginosa* isolates were removed from the 110 based on contamination, poor assembly completeness, and less than 10 SNP differences (identified using snp-dist (https://github.com/tseemann/snp-dists; Supplementary Table 1)). Isolate PAE36 had consistently poor growth and therefore, was removed. M1C124 and M1C010 were both kept despite their similarity because they were the only in-house representatives of the newly formed species, *Pseudomonas paraeruginosa*. This quality control left a panel of 88 *P. aeruginosa* that were used in subsequent analyses.

For pangenomic analyses, reference *P. aeruginosa* genomes from Ozer et. al., 2019 were downloaded using NCBI datasets v16.4.4 (16). The pangenome was determined with ppanggolin v2.0.5 and a phylogenetic tree created from the pangenome multiple sequence alignment using IQtree v2.0.3 and the JTT+F+I+G4 model with 1,000 ultra-fast bootstraps (17). The phylogenetic tree was visualised and annotated with ITOL (18) and where necessary, was supplemented with data from Ozer et. al., 2019 (19). Both the count and presence–absence (binary) matrices of defence system subfamilies were extracted from the PADLOC output files for each bacterial genome, except for M1C124 and M1C010 which were excluded. These matrices were then used to characterise the overall defence repertoire per isolate and to assess the prevalence of each subfamily across the panel. The distribution of defence systems per genome were visualised using a histogram and summarised by subfamily frequencies using a proportional area chart, aggregating rare systems into an "Other" category.

**Bacteriophage isolation, visualisation and Noongar descriptive naming**

Phages were isolated from wastewater collected at the Subiaco Wastewater Treatment Plant (Shenton Park, Perth, Western Australia) as previously described (20, 21). Phages were propagated and titred using standard double overlay agar methods (20, 21). Aliquots were prepared and visualised by transmission electron microscopy (TEM) by Mr Christopher Leigh (University of Adelaide, South Australia, Australia) as described (22). Since phages were isolated on the traditional homelands of the Noongar people, (Noongar Wadjuk Boodjar) candidates were assigned a descriptive nomenclature via consultation with the Noongar Language Centre (Perth, Western Australia, Australia) (Supplementary Table 2; Supplementary Figure 1).

**Bacteriophage DNA Extraction, whole genome sequencing and bioinformatic analyses**

Phage DNA was extracted from phage preparations as previously described (20, 21, 23). Whole genome sequencing of phage isolate DNA was performed by the Australian Genome Research Facility (AGRF; Melbourne, Victoria, Australia). Phage DNA libraries were prepared using the Nextera XT kit (Illumina Inc., USA) and sequenced on the NovaSeq6000 Illumina platform which produced 150bp paired-end reads. Genome assembly and putative phage contig identification were performed using the Phanta v.0.3 pipeline (24). To investigate the anti-defence repertoire of phages in our panel, we employed two complementary tools: DefenseFinder (v1.0.10) and dbAPIs (commit ID: 74af7e2) (25, 26). Both the HMMScan and DIAMOND outputs from dbAPIs were analysed. Final high confidence annotations were obtained by filtering for those with conditional e-value $\leq$ 1e−5 from the HMMScan output or e-value $\leq$ 1e−5 in the DIAMOND output. The final high confidence annotation was created by merging each of the search methods annotations, preferentially using the HMMScan result over the DIAMOND annotation. Annotations were labelled with the experimentally validated anti-defence seed protein whether it was annotated as part of the same family or clan. Some phage genes were annotated in a CLAN that contained both *tad1* and *tad2* and were labelled as *tad1* & *tad2*-like to differentiate these from genes that were within the same family as the *tad1* experimentally validated seed protein. The dbAPIs and DefenseFinder annotations were merged to achieve a consensus with preference given to the DefenseFinder annotations where there were conflicting results, given its stronger empirical basis.

Phage RBPs were predicted using the PhageRBPdetection v.4 pipeline (27). To focus on proteins plausibly involved in adsorption or structural interactions, the RBP matrix was filtered to exclude annotations corresponding to known or predicted anti-defence, or lytic effectors (e.g., anti-CRISPR Cas, CBASS, Gabija, Retron, Thoeris, endolysin). RBP and anti-defence annotations were aggregated for each phage, and a binary presence–absence matrix was constructed to represent the predicted RBP and anti-defence repertoire of each genome.

**Bacteriophage resistant mutant generation, sequencing and bioinformatic analyses**

A PAO1 mutant resistant to a *Phikzvirus* phage, J5TC, was generated and purified as previously described (28). DNA was extracted using the Qiagen DNeasy Tissue and Blood extraction kit, following manufacturer's instructions (Qiagen, Hilden, Germany). Bacterial DNA sequencing was performed by the Australian Genome Research Facility (AGRF; Melbourne, Victoria, Australia) as for the Illumina short read sequencing performed for the phages previously. The reads were compared to the PAO1 wild-type via Snippy v4.6.0 (29).

**Bacteriophage receptor efficiency of plating (EOP)**

Phage receptors were determined using a modified EOP as previously described (30). Ten-fold serial dilutions of phages were spotted onto the surface of overlay agar plates containing the isogenic PAO1 mutants or in-house PAO1 mutant, before incubation, titre calculation and EOP determination. EOPs were interpreted as previously described (31).

**Bacteriophage minimum lytic concentration (MLC)**

MLC of phages against *P. aeruginosa* isolates were determined as described with minor modifications (32). Briefly, phages were diluted to $10^8$ and $10^6$ plaque forming units per milliltre. Overlay agar plates were prepared by mixing 100-200 µL of each of the 88 *P. aeruginosa* isolates with 3 mL of overlay agar and pouring onto LB agar plates (Becton Dickinson, USA). Then, 5 µL of the phage dilutions were spotted onto the surface in duplicate. The plates were incubated at 37˚C overnight and inspected for signs of lysis. An MLC score for each phage-bacterial combination was given based on the amount of lysis (single plaques vs confluent lysis) at the lowest phage concentration as described (32).

**Relationship between host-range and phage genus and isolation host**

Phage–bacteria interaction profiles were binarised from MLC scores (lytic = 1 when MLC > 1; non-lytic = 0 when MLC ≤1), and pairwise Jaccard distances were calculated between phages based on lytic activity across the 88 *P. aeruginosa* isolates. Associations between host range, structure and either phage genus or isolation host were tested using permutational multivariate analysis of variance (PERMANOVA;999 permutations), and adjusted $R^2$ values were calculated from the pseudo-F statistic. Differences in permutational multivariate dispersion (PERMDISP) were evaluated to assess potential confounding. All analyses were conducted in Python 3.12.0 using scikit-bio v0.6.3 and associated packages (33).

#### Bacteriophage activity determinant association with activity

The filtered RBP set was used to compute per-phage RBP diversity metrics, including RBP richness (number of distinct RBP types) and Shannon entropy, calculated as:

$$H = \sum_{i=1}^{n} p_i \log_2 p_i$$

where $p_i$ is the proportion of the $i^{th}$ RBP type in the phage's repertoire.

To quantify the relationship between RBP diversity and host range, Spearman correlation tests were performed between RBP Shannon entropy and infectivity (percentage of susceptible bacterial strains). Pairwise Jaccard distances were also computed on the filtered RBP matrix and compared to host range dissimilarities using a Mantel test (Pearson correlation, 999 permutations). A complementary Spearman correlation was also computed between the two dissimilarity matrices. Finally, individual RBPs were ranked by their Spearman correlation with infectivity across the phage panel. The top 15 RBPs were plotted to assess the distribution of correlation coefficients and structural annotations (e.g., Tail, Head and Packaging, or Unannotated), enabling further insights into the functional contributions of specific RBP types.

#### *P. aeruginosa* defence systems and their influence on phage infectivity

To evaluate the influence of defence systems on phage infectivity, analyses were performed at the level of bacterial isolates using descriptive visualisations and mixed-effects modelling. First, defence system burden was related to phage infectivity using two complementary outcome measures: (i) the number of successful phage infections per isolate, defined as the number of phages exhibiting lytic activity (MLC > 1), and (ii) the average MLC across phages per isolate, restricting analyses to lytic interactions. These relationships were assessed using linear mixed-effects models with the number of defence systems as a fixed effect and bacterial phylogroup included as a random intercept to account for evolutionary structure. Model coefficients (β) and associated p-values were estimated by maximum likelihood, and fitted regression lines with 95% confidence intervals were overlaid on scatter plots. Next, defence system-specific patterns in susceptibility were examined by summarising phage infectivity for each defence system as the mean MLC across all tested phages for isolates encoding that system. The distribution of isolate-level mean MLC values was visualised using point-range plots showing the mean, minimum and maximum MLC across isolates. Associations between individual defence systems and resistance were assessed using one-sided Wilcox rank-sum tests comparing isolate-level mean MLC values between isolates with and without each system, restricted to defence systems detected in at least eight isolates.

**Univariate analysis of infectivity-associated features**

We screened all previously identified genomic features for association with phage infectivity using univariate tests across all phage-bacterium pairs. For count-based features, we compared MLC values between genomes with and without the feature using the Mann-Whitney U test. For continuous features, Spearman correlations were computed. Effect direction (ΔMLC or correlation coefficient) and p-values were recorded. Features present and absent in fewer than 10 phage-*P. aeruginosa* interactions were excluded. Each feature was annotated by functional class (RBP, defence, anti-defence), factors were considered significant if $p<0.05$ and were visualised in a volcano plot wherein point size reflected prevalence and point colour denoted feature class. This approach identified candidate features with strong individual influence on host-range, guiding subsequent multivariate modelling.

**Informed multivariate modelling of infectivity**

To assess the predictive utility of molecular features associated with infectivity, we trained a CatBoost Machine Learning (ML) classifier using features filtered through prior univariate analysis ($p < 0.05$; ≥10 genomes per class). Model performance was evaluated over 1000 stratified train-test splits (85:15), using fixed hyperparameters and random seeds for bootstrap resampling. At each iteration, Receiver operating characteristic (ROC)-area under the curve (AUC) scores, precision-recall curves, and confusion matrices were recorded. Mean AUC and 95% empirical confidence intervals were computed, and aggregated ROC and PR curves were generated with bootstrapped confidence bands to assess model robustness and stability.

7. Parks DH, Imelfort M, Skennerton CT, Hugenholtz P, Tyson GW. CheckM: assessing the quality of microbial genomes recovered from isolates, single cells, and metagenomes. Genome research. 2015;25(7):1043-55.
8. Li H. Minimap2: pairwise alignment for nucleotide sequences. Bioinformatics. 2018;34(18):3094-100.
9. Mikheenko A, Saveliev V, Gurevich A. MetaQUAST: evaluation of metagenome assemblies. Bioinformatics. 2015;32(7):1088-90.
10. Okonechnikov K, Conesa A, García-Alcalde F. Qualimap 2: advanced multi-sample quality control for high-throughput sequencing data. Bioinformatics. 2016;32(2):292-4.
11. Schwengers O, Jelonek L, Dieckmann MA, Beyvers S, Blom J, Goesmann A. Bakta: rapid and standardized annotation of bacterial genomes via alignment-free sequence identification. Microb Genom. 2021;7(11).
12. Seemann T. mlst. Github 2022.
13. Akhter S, Aziz RK, Edwards RA. PhiSpy: a novel algorithm for finding prophages in bacterial genomes that combines similarity- and composition-based strategies. Nucleic Acids Res. 2012;40(16):e126.
14. Payne LJ, Todeschini TC, Wu Y, Perry BJ, Ronson Clive W, Fineran Peter C, et al. Identification and classification of antiviral defence systems in bacteria and archaea with PADLOC reveals new system types. Nucleic Acids Res. 2021;49(19):10868-78.
15. Thrane SW, Taylor VL, Lund O, Lam JS, Jelsbak L. Application of Whole-Genome Sequencing Data for O-Specific Antigen Analysis and In Silico Serotyping of Pseudomonas aeruginosa Isolates. J Clin Microbiol. 2016;54(7):1782-8.
16. O'Leary NA, Cox E, Holmes JB, Anderson WR, Falk R, Hem V, et al. Exploring and retrieving sequence and metadata for species across the tree of life with NCBI Datasets. Scientific Data. 2024;11(1):732.
17. Gautreau G, Bazin A, Gachet M, Planel R, Burlot L, Dubois M, et al. PPanGGOLiN: Depicting microbial diversity via a partitioned pangenome graph. PLoS Comput Biol. 2020;16(3):e1007732.
18. Letunic I, Bork P. Interactive Tree Of Life (iTOL) v5: an online tool for phylogenetic tree display and annotation. Nucleic Acids Res. 2021;49(W1):W293-W6.
19. Ozer EA, Nnah E, Didelot X, Whitaker RJ, Hauser AR. The Population Structure of Pseudomonas aeruginosa Is Characterized by Genetic Isolation of exoU+ and exoS+ Lineages. Genome biology and evolution. 2019;11(1):1780-96.
20. Vaitekenas A, Tai AS, Ramsay JP, Stick SM, Agudelo-Romero P, Kicic A. Complete Genome Sequences of Four Pseudomonas aeruginosa Bacteriophages: Kara-mokiny 8, Kara-mokiny 13, Kara-mokiny 16, and Boorn-mokiny 1. Microbiology Resource Announcements. 2022;11(12):e00960-22.
21. Ng RN, Tai AS, Chang BJ, Stick SM, Agudelo-Romero P, Kicic A. Complete Genomes of Three Pseudomonas aeruginosa Bacteriophages, Kara-mokiny 1, Kara-mokiny 2, and Kara-mokiny 3. Microbiology Resource Announcements. 2022;11(12):e00955-22.
22. Iszatt JJ, Larcombe AN, Garratt LW, Stick SM, Kicic A. Lytic activity, stability, biofilm disruption capabilities, and genomic characterization of two bacteriophages active against respiratory MRSA. Journal of Applied Microbiology. 2025;136(4).

23. Jakočiūnė D, Moodley A. A Rapid Bacteriophage DNA Extraction Method. Methods and protocols. 2018;1(3).
24. Iszatt J. Phanatic. 0.3 ed: Github; 2023.
25. Yan Y, Zheng J, Zhang X, Yin Y. dbAPIS: a database of anti-prokaryotic immune system genes. Nucleic Acids Res. 2023;52(D1):D419-D25.
26. Tesson F, Huiting E, Wei L, Ren J, Johnson M, Planel R, et al. Exploring the diversity of anti-defense systems across prokaryotes, phages and mobile genetic elements. Nucleic Acids Res. 2024;53(1).
27. Boeckaerts D, Stock M, De Baets B, Briers Y. Identification of Phage Receptor-Binding Protein Sequences with Hidden Markov Models and an Extreme Gradient Boosting Classifier. Viruses. 2022;14(6).
28. Wannasrichan W, Htoo HH, Suwansaeng R, Pogliano J, Nonejuie P, Chaikeeratisak V. Phage-resistant Pseudomonas aeruginosa against a novel lytic phage JJ01 exhibits hypersensitivity to colistin and reduces biofilm production. Frontiers in Microbiology. 2022;13.
29. Seeman T. Snippy. 4.6.0 ed: Guthub; 2020.
30. Khan Mirzaei M, Nilsson AS. Isolation of phages for phage therapy: a comparison of spot tests and efficiency of plating analyses for determination of host range and efficacy. PLoS One. 2015;10(3):e0118557.
31. Gibson SB, Green SI, Liu CG, Salazar KC, Clark JR, Terwilliger AL, et al. Constructing and Characterizing Bacteriophage Libraries for Phage Therapy of Human Infections. Frontiers in Microbiology. 2019;10(2537).
32. Gaborieau B, Vaysset H, Tesson F, Charachon I, Dib N, Bernier J, et al. Prediction of strain level phage–host interactions across the Escherichia genus using only genomic information. Nature Microbiology. 2024;9(11):2847-61.
33. Jai Ram Rideout EB, Daniel McDonald, Yoshiki Vázquez Baeza, Jorge Cañardo Alastuey, Anders Pitman, Jamie Morton, Qiyun Zhu, Jose Navas, Kestrel Gorlick, Matt Aton, Justine Debelius, Zech Xu, llcooljohn, Joshua Shorenstein, Laurent Luce, Will Van Treuren, John Chase, charudatta-navare, Antonio Gonzalez, Colin J. Brislawn, Weronika Patena, Karen Schwarzberg, teravest, Igor Sfiligoi, Jens Reeder, Greg Caporaso, shiffer, nbresnick and Chris Tapo. scikit-bio. 0.7.0 ed: Zenodo; 2025.
